# Oleic acid triggers hippocampal neurogenesis by binding to TLX/NR2E1

**DOI:** 10.1101/2020.10.28.359810

**Authors:** Prasanna Kandel, Fatih Semerci, Aleksandar Bajic, Dodge Baluya, LiHua Ma, Kevin Chen, Austin Cao, Tipwarin Phongmekhin, Nick Matinyan, William Choi, Alba Jiménez-Panizo, Srinivas Chamakuri, Idris O. Raji, Lyra Chang, Pablo Fuentes-Prior, Kevin R. MacKenzie, Caroline L. Benn, Eva Estébanez-Perpiñá, Koen Venken, David D. Moore, Damian W. Young, Mirjana Maletic-Savatic

## Abstract

Adult hippocampal neurogenesis underpins learning, memory, and mood, but diminishes with age and illness. The orphan nuclear receptor TLX/NR2E1 is known to regulate neural stem and progenitor cell self-renewal and proliferation, but the precise mechanism by which it accomplishes this is unknown. We found that neural stem and progenitor cells require monounsaturated fatty acids to survive and proliferate. Specifically, oleic acid (18:1ω9) binds to TLX to convert it from a transcriptional repressor to a transcriptional activator of cell cycle and neurogenesis genes. We propose a model in which sufficient quantities of this endogenous ligand must bind to TLX to trigger the switch to proliferation. These findings pave the way for future therapeutic manipulations to counteract pathogenic impairments of neurogenesis.

---

The mammalian brain contains a population of neural stem cells that continue to produce functional neurons well into adulthood (*1–4*). This ongoing capacity to form new neurons, known as adult neurogenesis, enables learning and memory (*5, 6*) and supports proper mood regulation (*7*). Conversely, impairments in neural stem cell proliferative capacity and neuronal commitment are associated with advanced age (*8, 9*) and disorders such as depression (*10*) and Alzheimer’s disease *(11, 12).* Indeed, antidepressant therapies such as serotonin reuptake inhibitors are effective only insofar as they succeed in stimulating neurogenesis (*7*). There has thus been considerable interest in learning how to manipulate hippocampal neurogenesis in order to treat a range of conditions.

One key question is how to preserve the pool of neural stem and progenitor cells while promoting the ongoing generation of neurogenic progeny. To maintain their population, these cells must be able to self-renew, and it is well established that neural stem and progenitor cell self-renewal and proliferation are regulated by the transcription factor TLX (also known as NR2E1) (*13–15*). Unfortunately, TLX is an orphan nuclear receptor—in other words, no endogenous ligand for TLX has yet been discovered (exogenous and synthetic ligands have been reported but without physiological relevance *in vivo* (*16, 17*)). Without such a ligand, TLX cannot be validated as a therapeutic target for promoting neurogenesis.

We are not without clues as to the probable nature of such a ligand, however. First, nuclear receptors as a class tend to bind to lipophilic molecules, such as hormones, vitamins, and fatty acids. Second, neural stem cells require *de novo* lipogenesis to proliferate (*18–20*). Third, we previously identified a nuclear magnetic resonance (NMR) signal that correlates with neurogenic activity in mice and humans; it resonates at 1.28 ppm, a frequency that corresponds to the saturated hydrocarbon groups that characterize all fatty acids (*21*). Fourth, blocking the activity of fatty acid synthase, which catalyzes the synthesis of saturated fatty acids, markedly reduces the 1.28 ppm NMR signal (*21*) and prevents running from stimulating neurogenesis in mice (*18*). We therefore hypothesized that fatty acids would be crucial to TLX function.

In fact, compared to other cells such as astrocytes and fully differentiated neurons, neural stem cells have particularly high levels of mono-unsaturated fatty acids (MUFAs) (*21*). Interestingly, while most of the metabolic genes enriched in quiescent neural stem cells are associated with fatty acid synthesis, these cells have particularly high expression of stearoyl-CoA desaturases (*22, 23*), which are involved specifically in the synthesis of MUFAs. This suggests that MUFAs might have a special role in controlling the cell cycle in neural stem cells. In this study, we asked whether MUFA synthesis is necessary for survival and proliferation of these cells, and whether these fatty acids might act as ligands for TLX in neural stem cells.

## MUFAs are essential for neural stem and progenitor cell survival and proliferation

To begin to address these questions, we first confirmed that MUFAs are abundant in human neural stem and progenitor cells. Two-dimensional ^1^H-^1^H COSY NMR, which identifies coupled hydrogen (^1^H) spins, revealed the mono-unsaturated bond of MUFAs in these cells (fig. S1A). Next, gas chromatography–mass spectrometry found several saturated fatty acid precursors and MUFAs, with oleic acid (18:1ω9) being the most abundant (fig. S1B). Like their murine counterparts, human neural stem and progenitor cells require fatty acid synthase to survive: cerulenin, an inhibitor of fatty acid synthase, decreased their viability in a dose-dependent manner (fig. S2A). Treating the cells with CAY10566, an inhibitor of the stearoyl-CoA desaturases that catalyze the conversion of saturated fatty acids into MUFAs, resulted in a dose- and time-dependent impairment of survival *in vitro* (Fig. 1A, left panel). Strikingly, this impairment could be rescued by exogenous 18:1 MUFA but not 18:0 saturated fatty acids (Fig. 1A, right panel; fig. S2B). Consistent with these *in vitro* data, oral administration of CAY10566 in mice significantly reduced neural stem and progenitor cell proliferation *in vivo* (Fig. 1B). Therefore, human neural stem and progenitor cells require *de novo* MUFA synthesis for proliferation and survival.

**Fig. 1.**
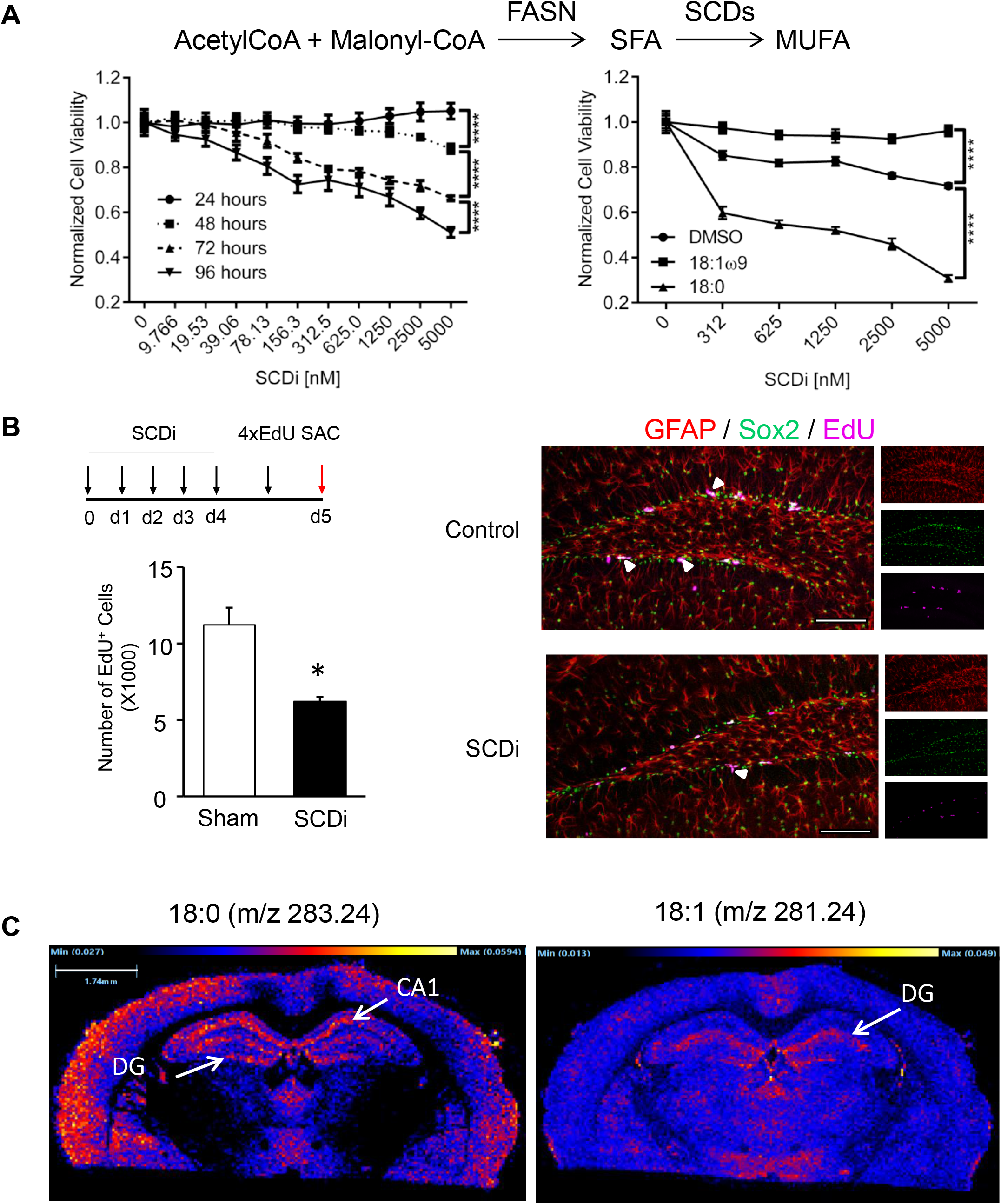
Human neural stem and progenitor cells depend on *de novo* mono-unsaturated fatty acid (MUFA) synthesis to survive and proliferate. **(A)** Acetyl coenzyme A (AcetylCoA) and methyl-malonyl CoA serve as substrates for saturated fatty acid (SFA) synthesis catalyzed by fatty acid synthase (FASN). SFAs are converted to MUFAs via desaturases, such as the stearoyl-CoA desaturases (SCD), which catalyze the rate-limiting step in the MUFA synthesis. *Left panel:* Dose-dependent human neural stem and progenitor cell viability following SCD inhibition with CAY10566 (SCDi), normalized to vehicle (DMSO) response. *Right panel:* Dose-dependent human neural stem and progenitor cell viability following 144 hr exposure to SCDi and then treated with 18:1ω9 MUFA or its precursor, 18:0 SFA, normalized to the respective treatment (N≥3 per group). Statistics was done using two-way ANOVA and Tukey’s multiple comparisons test, ****p≤0.0001. **(B)** SCDi (3 mg/kg body weight) or sham (0.03N HCl) was delivered orally to 2-3-month-old wild-type C57BL/6J mice followed by EdU (50 mg/kg body weight) to label proliferating human neural stem and progenitor cells *in vivo* (N=3 per group). Bar graphs represent mean ± SEM. *p≤0.05. Representative confocal micrographs show proliferating Edu^+^ GFAP^+^ Sox2^+^ human neural stem and progenitor cells. Scale bar=100 μm. **(C)** Imaging mass spectrometry of the 3-month-old wild-type C57BL/6J mouse brain shows distribution of the 18:0 SFAs (m/z 283.24, *left panel*), the precursor of 18:1 MUFAs (m/z 281.24, *right panel*). Dentate gyrus (DG) and CA1 region of hippocampus are indicated (arrows). See also figs. S1-3.

If MUFA synthesis is required for survival of these cells, then MUFAs should be abundant in the dentate gyrus of the hippocampus, where neurogenesis occurs. Using imaging mass spectrometry to map lipids in the mouse brain, we detected both 18:0 saturated fatty acids (*m*/*z*=283.24) and 18:1 MUFA (*m*/*z*=281.24) in the dentate gyrus. In order to gauge the concentration, we spiked 0.5 mM 18:1ω9 control onto the thalamus (Fig. 1C, fig. S3). The 18:1 MUFA ion showed a very similar concentration in the dentate gyrus, where it was most abundant. We therefore considered the possibility that the MUFAs are not being used just for energy metabolism, lipid membrane formation, and so forth, but could also serve as signaling molecules that regulate neurogenesis.

## Oleic acid binds to the TLX ligand-binding domain

Because there is no crystal structure of TLX bound to ligands, we searched for a nuclear receptor that binds fatty acids and whose ligand-binding domain has an established structure. The closest liganded homolog is HNF4α (fig. S4). Initial structural studies revealed HNF4α bound to fatty acids (*24, 25*) and functional studies indicate that HNF4α is selectively activated by exogenous linoleic acid (C18:2ω6) in mammalian cells and in the liver of fed mice (*26*). We therefore built a homology model of the human TLX ligand-binding domain and performed molecular docking based on the X-ray crystal structure of fatty acid-bound HNF4α ligand-binding domain (PDB 1M7W). These docking studies suggested that fatty acids could indeed fit into the binding pocket and potentially function as cognate ligands of TLX (fig. S4E).

To determine whether fatty acids actually bind to the purified TLX ligand-binding domain (fig. S5A), we used biolayer interferometry to compare the binding responses of different fatty acids with an18 carbon chain length (Fig. 2A). Stearic acid (18:0 saturated fatty acid) had no observable binding, while α-linolenic acid (18:3ω3 poly-unsaturated fatty acid) failed to reach equilibrium or saturate over the concentrations tested, suggesting a weak binding interaction. Oleic acid (18:1ω9 MUFA), however, displayed a saturable binding response with a K_D_ of 7.3 μM. Interestingly, the synthetic 18:1ω5 MUFA had a similar saturable binding response with a K_D_ of 6.5 μM (fig. S5B). These experiments show that MUFAs, such as 18:1ω9, can bind to the TLX ligand-binding domain.

**Fig. 2.**
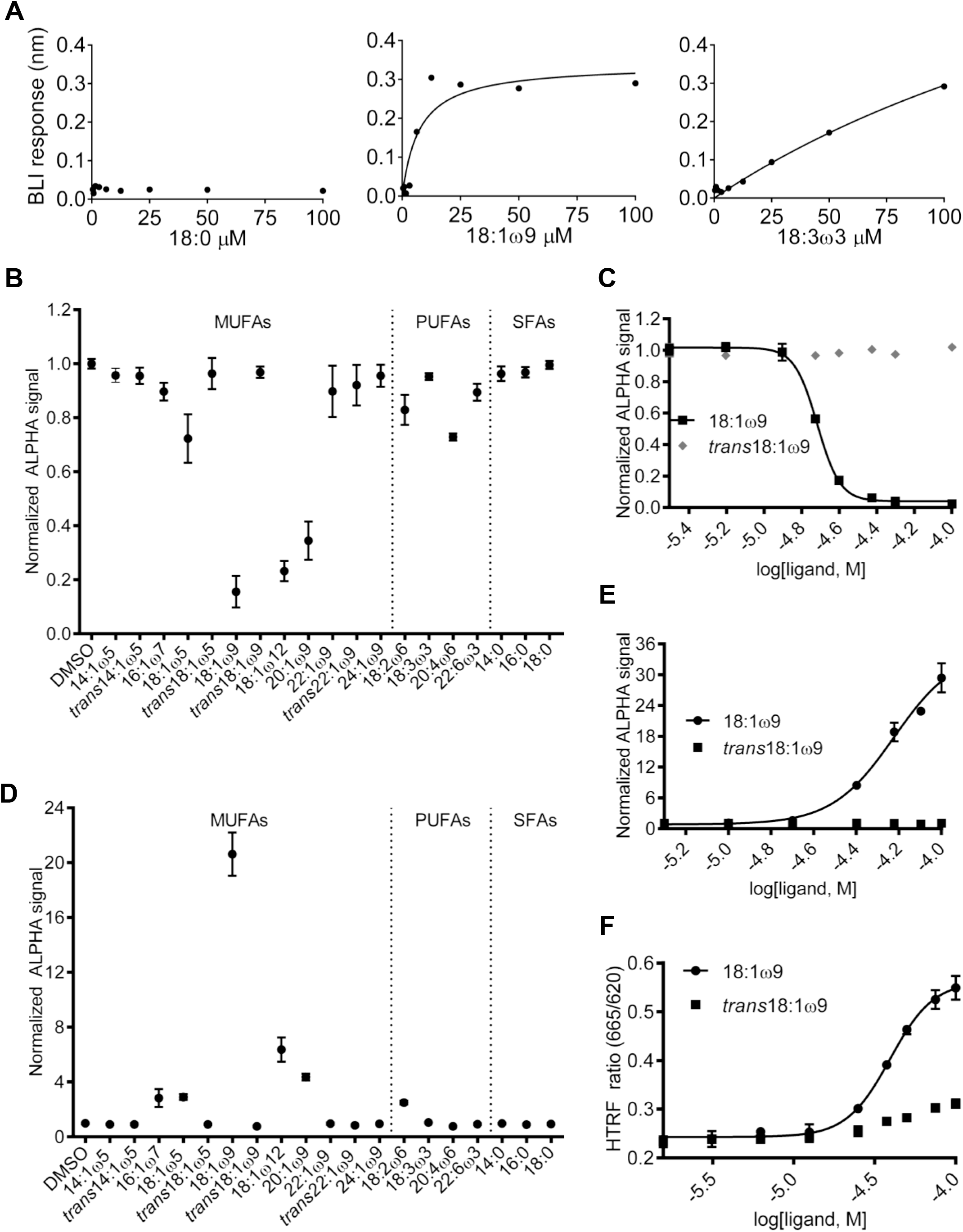
18:1ω9 MUFA binds to TLX ligand-binding domain, reduces the corepressor and enhances the coactivator interaction. **(A)** Dose-dependent binding response of different classes of fatty acids with the same carbon chain length (18 carbons) to the TLX ligand-binding domain, using biolayer interferometry label-free technology. 18:0 SFA has no observable binding response (*left panel*). 18:1ω9 MUFA binding response fit is a saturable one-site fit (the dissociation constant (K_d_) =7.3 μM, 95% CI=0.385-14.2 μM; *middle panel*). 18:3ω3 polyunsaturated fatty acid (PUFA) binding response fit is non-saturable (K_d_=270.9 μM, 95% CI=140.8-400.9 μM, *right panel*). **(B)** Normalized Amplified Luminescent Proximity Homogeneous Assay (ALPHA) screen for the interaction between TLX ligand-binding domain and a corepressor (atrophin peptide) in the presence of selected MUFAs, PUFAs, and SFAs at 25 μM. Signal <1.0 indicates decreased interaction. **(C)** Dose-dependent TLX ligand-binding domain/corepressor interaction in the presence of 18:1ω9 (EC_50_=19.3 μM, 95% CI=18.7-19.5 μM) or *trans*18:1ω9 MUFA (synthetic stereoisomer). **(D)** Normalized ALPHA screen of the interaction between TLX ligand-binding domain and a coactivator (NCOA1 peptide) in the presence of selected MUFAs, PUFAs, and SFAs screened at 50 μM. Signal >1.0 indicates increased interaction. Note the interaction in the presence of 18:1ω9 versus *trans*18:1ω9. **(E)** Dose-dependent TLX ligand-binding domain/coactivator interaction in the presence of 18:1ω9 (EC_50_=59.8 μM, 95% CI=50.6-70.8 μM) or *trans*18:1ω9. **(F)** Dose-dependent Homogeneous Time Resolved Fluorescence (HTRF) signal for TLX/coactivator peptide recruitment in the presence of *cis* vs. *trans* 18:1 MUFA (EC_50_=39.3 μM for 18:1ω9, 95% CI=37.0-41.7 μM). See also figs. S4-6.

## Oleic acid binding to TLX promotes recruitment of co-activators

To understand the functional effects of MUFA binding to TLX, we considered that in its basal state, TLX recruits corepressor proteins such as atrophins (ATN 1 and 2) (*27*), lysine-specific demethylase (LSD1) (*28*), and histone deacetylases (HDAC 1, 3, and 5) to potently repress transcription (*29*). We first confirmed the direct interaction between TLX and the atrophin corepressor peptide using biolayer interferometry and found that the atrophin peptide binds TLX with a K_D_ of 14.2 μM (fig. S6A). We next used the Amplified Luminescent Proximity Homogeneous Assay (ALPHA) to measure the interaction of atrophin peptide and TLX ligand-binding domain. Competition with untagged atrophin peptide attenuated the ALPHA signal with an EC_50_ of 1.3 μM, demonstrating that the assay can detect both TLX-corepressor binding and its disruption (fig. S6B). To determine which fatty acids could disrupt TLX-atrophin peptide interaction, we examined a panel of fatty acids (saturated, mono-unsaturated, and poly-unsaturated) (Fig. 2B). The saturated fatty acids (14:0, 16:0 and 18:0) had no effect on the ALPHA signal, indicating that they do not interfere with the interaction of atrophin with the TLX ligand-binding domain. The poly-unsaturated fatty acids (18:2ω6 and 20:4ω6) were, at best, only modestly active. Among the MUFAs, both the short- and long-chain fatty acids (e.g., 14:1ω5 and 24:1ω9) were inactive. The medium-chain MUFAs, however—18:1ω9 (oleic acid), 18:1ω12, and 20:1ω9—clearly disrupted the interaction of TLX with the corepressor. Out of the entire panel, *cis* 18:1ω9 (oleic acid) reduced the interaction with atrophin the most. Given that it was also the most abundant MUFA in human neural stem and progenitor cells (fig. S1B) and the most potent fatty acid in the panel at a single dose, we next performed dose-response experiments. These confirmed that 18:1ω9, but not its *trans* isomer, disrupts the interaction of the TLX ligand-binding domain with the atrophin peptide in a dose-dependent manner, with an EC_50_=19.3 μM (Fig. 2C). We conclude that the TLX ligand-binding domain stereospecifically recognizes certain fatty acids and that occupancy of the TLX ligand-binding pocket by specific MUFAs, especially oleic acid, elicits conformational changes that disengage the atrophin corepressor.

Could fatty acid ligands induce TLX to switch from corepressor to coactivator binding? TLX interacts with nuclear receptor coactivators such as NCOA1, NCOA2, NCOA3 (SRC1-3) (*30*). To examine whether oleic acid-bound TLX could recruit NCOA1, we again used the ALPHA screen, placing a tagged NCOA1-II coactivator peptide containing the canonical LXXLL nuclear receptor interaction motif and the TLX ligand-binding domain on donor and acceptor beads, respectively. In the absence of fatty acids or in the presence of only saturated or poly-unsaturated fatty acids, there was no interaction between the NCOA1-II peptide and the TLX ligand-binding domain, but 18:1ω9 markedly increased the ALPHA signal (Fig. 2D). As with disruption of the TLX-corepressor interaction, enhancement of the TLX-coactivator interaction was specific to 18:1ω9 and not *trans*18:1ω9, confirming dose-dependent stereospecificity of fatty acid binding and coactivator recruitment (Fig. 2E). To determine the affinity of coactivator binding, we performed competition experiments and found that, in the presence of saturating 18:1ω9, TLX recruited both the NCOA1-II peptide and a receptor-interaction domain from NCOA3, with EC_50_ values of 6.0 μM (fig. S6C) and 3.3 μM (fig. S6D), respectively. A TLX-based Homogeneous Time-Resolved Fluorescence assay independently verified ligand-dependent coactivator recruitment and the specificity of *cis/trans*18:1ω9 recognition (Fig. 2F). These data strongly indicate that oleic acid binding determines whether TLX functions as a transcriptional repressor or an activator.

To examine the effect of 18:1ω9 on TLX-mediated transcription, we built a dual luciferase transcription reporter system with a previously reported TLX-binding response element (*31*). We cotransfected TLX with the reporter plasmids in HeLa cells grown in media with charcoal-stripped fetal bovine serum to minimize fatty acids and other non-polar molecules. Luciferase expression was reduced in these cells compared to cotransfection with the empty TLX expression vector (Fig. 3A, left panel), confirming the repressive function of TLX at baseline. The predicted agonist activity of 18:1ω9 was confirmed by the expected dose-dependent increase in luciferase expression (Fig. 3A, middle panel). This effect was not seen with *trans*18:1ω9 (Fig. 3A, right panel; fig. S6E), verifying that the modest and stereospecific recognition of the *cis*-form was retained in living cells and that 18:1ω9 binding promotes TLX-mediated transcriptional activation of TLX-dependent genes.

**Fig. 3.**
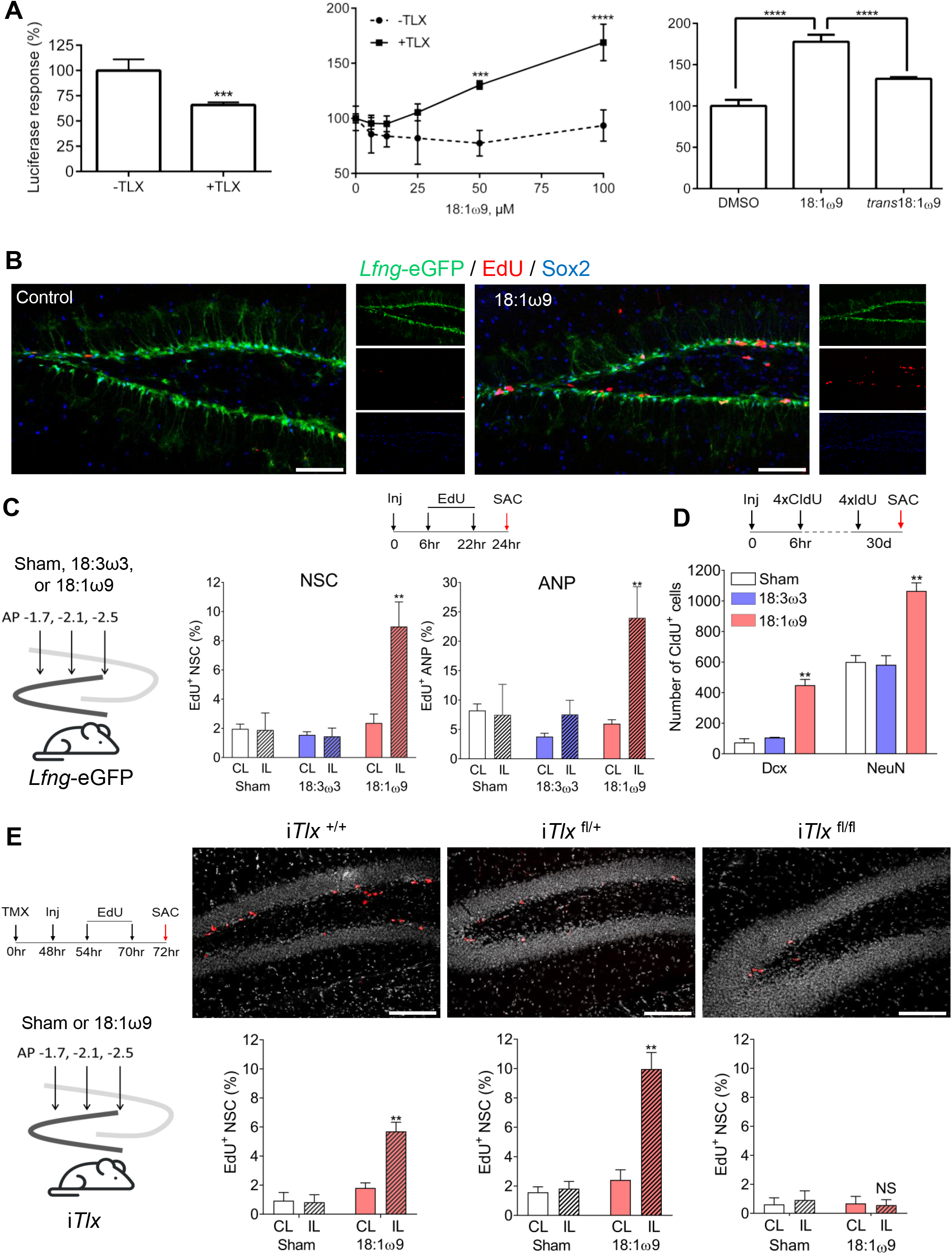
18:1ω9-dependent TLX effect on murine adult neural stem cell proliferation and neurogenesis *in vivo*. **(A)** TLX-based dual luciferase reporter activity in HeLa cells with or without TLX-expressing plasmids (*left panel*). Dose-dependent TLX-luciferase response in HeLa cells with or without TLX-expressing plasmids (*middle panel*). TLX-luciferase response in HeLa cells with TLX-expressing plasmids in the presence of *cis* or *trans* 18:1ω9 (100 μM each) (*right panel*). Bar graphs represent mean ± SEM. ***p≤0.005, **** p≤0.001. **(B)** Representative confocal hippocampal sections show *Lfng*-eGFP^+^ neural stem cells (NSCs), Sox2^+^ amplifying neuroprogenitors (ANPs), and EdU^+^ proliferating cells in the subgranular zone of a 2-month-old *Lfng*-eGFP mouse treated with either sham or 18:1ω9. Scale bar=100 μm. **(C)** The number of NSCs and ANPs in 2-month-old *Lfng*-eGFP mice injected with either sham (no solvent), 18:1ω9, or 18:3ω3 fatty acids into the left dentate gyrus (ipsilateral, IL) (300 nL of pure fatty acid per injection site); the right dentate gyrus (contralateral, CL) served as a non-injection control (N=3-5 per group). Sham served as an injection site control. 18:3ω3 served as a fatty acid control. Bar graphs represent mean ± SEM. For statistical analysis (two-way ANOVA, Tukey’s multiple comparison test), CL and IL from the same mouse were compared (Table 1A). **(D)** The number of newborn immature neurons and granule cells in 2-month-old *Lfng*-eGFP mice treated bilaterally with either sham (no solvent), 18:1ω9, or 18:3ω3 fatty acids and CldU (85 mg/kg) 6 hr thereafter (N=3-4 per group). Bar graphs represent mean ± SEM; statistics was done using one-way ANOVA and Tukey’s multiple comparison test (Table 2). Please see fig. S7 for quantification of IdU^+^ NSCs and ANPs 1-month post-injection. **(E)**2-month-old *Lfng*-CreER^T2^/Rosa26tdT/*Lfng*-eGFP/*Tlx*^fl/fl^ (i*Tlx*^fl/fl^) mice were given tamoxifen (TMX; 120 mg/kg body weight) to induce CreER^T2^ 48 hr before either sham (no solvent) or 18:1ω9 delivery (300 nL of pure fatty acid per injection site), followed by EdU (50 mg/kg i.p.) 6 hr and 22 hr thereafter (N=3 per group). Confocal micrographs show EdU^+^ proliferating cells in mice with two copies of *Tlx* (i*Tlx*^+/+^), one copy of *Tlx* (i*Tlx*^fl/+^), and lacking both copies of *Tlx* (i*Tlx*^fl/fl^). Bar graphs (mean ± SEM) show the percentage of proliferating EdU^+^ NSCs in i*Tlx*^+/+^, i*Tlx*^fl/+^, and i*Tlx*^fl/fl^ mice. For statistical analysis (two-way ANOVA, Tukey’s multiple comparison test), CL and IL from the same mouse were compared (Table 1B). *p≤0.05, ** p≤0.001, NS=non-significant. See fig. S8 for generation of i*Tlx*^fl/fl^ mice.

## Oleic acid-activated TLX triggers neural stem cell mitosis and neurogenesis *in vivo*

To test the effect of 18:1ω9 on adult neurogenesis *in vivo*, we stereotactically delivered either 18:1ω9, 18:3ω3 (as a fatty acid control), or sham to the left dentate gyrus, then treated mice with BrdU analogs to label newborn cells, and sacrificed them at different timepoints post-injection (Fig. 3B-D; Tables S1, S2). Compared to controls, 24 hr after injection the percentage of proliferating EdU^+^ neural stem cells and their progeny (amplifying neuroprogenitors) was elevated at the site of 18:1ω9 injection only (Fig. 3B, C; Table S1A). The stimulatory effect of 18:1ω9 is thus specific and local. The number of immature (Dcx^+^, CldU^+^) and mature (NeuN^+^, CldU^+^) newborn neurons rose 30 days following 18:1ω9 injection (Fig. 3D; Table S2), confirming neurogenesis had been stimulated. The percentage of proliferating (IdU^+^) neural stem cells and amplifying neuroprogenitors at this timepoint did not, however, differ from controls, indicating that the stimulatory effect of exogenous 18:1ω9 on these cells is short-lived (fig. S7).

To determine whether 18:1ω9 requires TLX to affect neural stem cell function, we took advantage of *Lfng*-eGFP transgenic mice, in which eGFP selectively labels radial neural stem cells (*32*). We crossed *Tlx*^loxp/loxp^ mice (*13*) with *Lfng*-eGFP;*Lfng-*CreER^T2^;RCL-tdT mice to achieve homozygous deletion of *Tlx* in Cre-induced neural stem cell following tamoxifen administration (fig. S8). In the resulting i*Tlx*^fl/fl^ mice, wild-type neural stem cells (no Cre activation) have two copies of *Tlx* (*Tlx*^+/+^) and are labeled green (eGFP). Mutant neural stem cells (Cre-induced) lack both copies of *Tlx* (*Tlx*^fl/fl^) and are labeled both red (tdT) and green (eGFP). Thus, in i*Tlx*^fl/fl^ brains we can examine 18:1ω9-dependent cell-autonomous effects in both wild-type and mutant neural stem cells from the same dentate gyrus. Following a single 18:1ω9 injection, the ratio of proliferating neural stem cells in i*Tlx*^fl/fl^ mice correlated with *Tlx* gene dosage: the percentage of EdU^+^ cells was much higher when two copies of *Tlx* were present than when only one copy was present, and 18:1ω9 exerted no proliferative effect when both copies of *Tlx* were lacking (Fig. 3E; Table S1B). Thus, independent of any purely metabolic role for MUFA, the stimulatory effects of 18:1ω9 on adult hippocampal neural stem cells and neurogenesis are mediated through TLX.

Our next step was to determine whether 18:1ω9-activated neural stem cell proliferation correlates with TLX target gene expression *in vivo*. We injected either 18:1ω9 or 18:3ω3 into both dentate gyri of *Lfng*-eGFP mice and performed RT-PCR on sorted neural stem cells to measure expression of known TLX targets as well as a panel of cell cycle and neurogenesis genes. To qualify as being activated in a 18:1ω9-dependent manner, a given gene should be upregulated ≥4-fold by 18:1ω9 and not affected by 18:3ω3. We found 92 such genes (55 involved in the cell cycle and 37 in neurogenesis) (Fig. 4A). (An additional 11 cell cycle and 8 neurogenesis genes were upregulated by both 18:1ω9 and 18:3ω3 [Fig. 4A; fig. S9A]; these may be nonspecific fatty acid responders and were excluded from further analyses.)

**Fig. 4.**
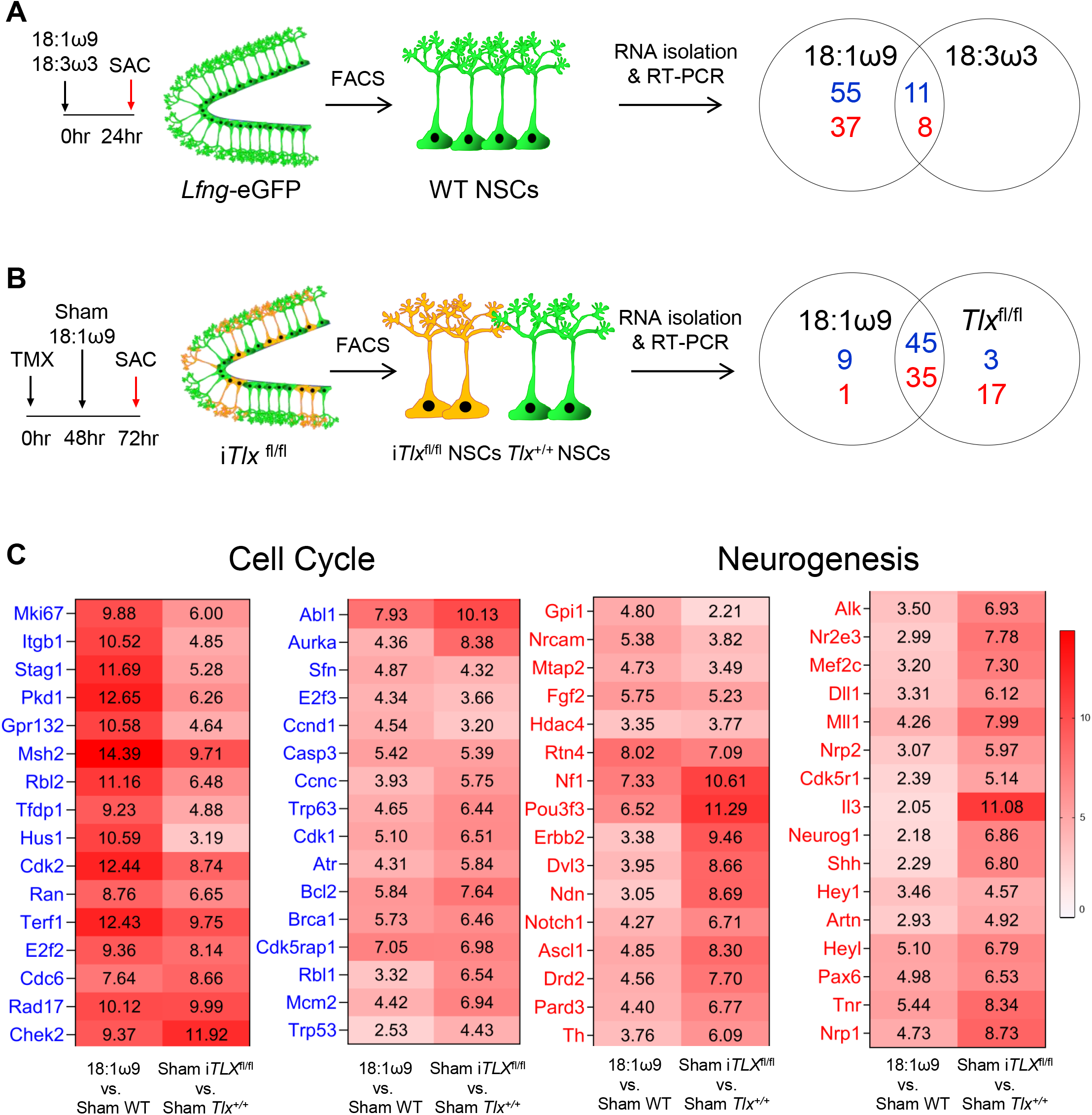
18:1ω9 induces TLX to upregulate cell cycle and neurogenesis genes in adult murine hippocampal neural stem cells in a TLX-dependent manner. **(A)** 2-month-old *Lfng*-eGFP mice—in which all NSCs are labeled green—were treated with either 18:1ω9, 18:3ω3 (fatty acid control), or sham (N=3 per group), and eGFP^+^ NSCs were sorted from the dentate gyri. RT-PCR was done using a panel of cell cycle and neurogenesis genes. Venn diagram represents differentially regulated genes derived from pairwise ΔC_t_ value comparison across groups. Numbers in blue are cell cycle genes and numbers in red are neurogenesis genes. **(B)** 2-month-old i*Tlx*^fl/fl^ (*Lfng*-CreER^T2^/Rosa26tdT/ *Lfng*-eGFP/*Tlx*^fl/fl^) mice were given tamoxifen (TMX; 120 mg/kg body weight) 48 hr before either sham (no solvent) or 18:1ω9 delivery (N=3 per group). *Tlx*^fl/fl^ NSCs (tdT^+^ eGFP^+^; orange) and *Tlx*^+/+^ NSCs (eGFP^+^) were sorted from the dentate gyri. RT-PCR was done for the same cell cycle and neurogenesis genes as in Fig. 4A. Venn diagram represents differentially regulated genes derived from pairwise ΔC_t_ value comparison across groups. Numbers in blue are cell cycle genes and numbers in red are neurogenesis genes. **(C)** Heatmaps of ΔΔC_t_ values (cited in boxes) show 32 cell cycle and 32 neurogenesis genes exclusively upregulated by 18:1ω9 (≥4-fold change) in a TLX-dependent manner. 18:1ω9-treated vs. sham-treated *Tlx*^+/+^ NSCs (*Lfng*-eGFP^+^ NSCs from *Lfng*-eGFP mice) indicates genes upregulated by 18:1ω9. Sham-treated *Tlx*^fl/fl^ NSCs (*Tlx*^fl/fl^ tdT^+^ eGFP^+^) vs. sham-treated *Tlx*^+/+^ NSCs indicates genes de-repressed by TLX. See also fig. S9.

To determine which of the 92 18:1ω9-dependent genes are also TLX-dependent, we injected either 18:1ω9 or sham into both dentate gyri of i*Tlx*^fl/fl^ mice and performed RT-PCR on sorted neural stem cells again, using the same gene panels (Fig. 4B). We identified 48 cell cycle and 52 neurogenesis genes upregulated ≥4-fold in neural stem cells lacking *Tlx* (*Tlx*^fl/fl^ eGFP^+^, tdT^+^) compared to wild-type neural stem cells (*Tlx*^+/+^ eGFP^+^) from sham-treated *iTlx*^fl/fl^ mice. Among these genes, 45 cell cycle and 35 neurogenesis genes overlapped with those upregulated by 18:1ω9 (Fig. 4B). Of those, 32 cell cycle and 32 neurogenesis genes were upregulated by 18:1ω9 in a TLX-dependent manner (Fig. 4C). We did not pursue the basis of the response of the remaining 13/45 cell cycle and 3/35 neurogenesis genes that were not exclusively TLX-dependent (fig. S9B, C), although they warrant further study.

The 18:1ω9-activated TLX-dependent genes included previously reported TLX targets such as *Ascl1* (*13*), *Trp53* (*33*), *Mcm2,* and *Ccnd2* (*34*), but also revealed new targets such as *Olig2, Shh, Fgf2, Dvl3, Hey1, Pax6, Nf1, Dll1*, *Nr2e3*, *E2f2, E2f3, Cdk1,* and *Cdk2*. It is worth noting that TLX does not act as a binary on-off switch: 18:1ω9-bound TLX activates cell cycle genes more strongly than neurogenesis genes, while lack of TLX de-represses neurogenesis genes more strongly (Fig. 4C). This is consistent with previous reports that a gradual increase—and not a steady, high expression—of neurogenic genes, such as *Ascl1*, is necessary for neuroprogenitors to slowly enter into cell cycle to avoid premature differentiation (*35–37*).

## Discussion

This study thus sheds new light on the importance of fatty acid metabolism and mono-unsaturated fatty acids as signaling molecules. Oleic acid is not only essential for neural stem cell proliferation and neurogenesis in the hippocampus, but it appears to help preserve the neural stem cell population over time. Because TLX activation requires relatively high levels of oleic acid, neural stem cells are not perpetually dividing and quickly depleting in number. Our results support a model in which oleic acid acts as a critical signaling metabolite in neural stem cells. In its ligand-free state, TLX favors corepressor binding and repression of its target genes, causing these cells to remain quiescent. During quiescence, oleic acid is synthesized and accumulates until it reaches a certain threshold that indicates the availability of metabolic prerequisites for cell cycle entry. Upon binding to oleic acid, TLX disengages from the corepressors and recruits transcriptional coactivators, which triggers neural stem cell proliferation. In response to acutely increased oleic acid levels, robust upregulation of cell cycle genes would prompt neural stem cells to generate neurogenic progeny while preventing their direct differentiation into neurons. Temporally cued de-repression of neurogenesis genes could then ensure that the progeny properly proceed into the neurogenic and not astrocytic lineage.

A recent study reported that TLX is also a receptor for exogenous natural and synthetic retinoids but did not explore the physiology of these interactions *in vivo* (*16*). Here, by demonstrating that oleic acid binding to TLX is crucial for stimulating neurogenesis, this work lays a foundation for future efforts to preserve neural stem cells during aging and in disease. The discovery of an endogenous ligand for TLX will facilitate efforts to identify small molecules that can modulate TLX function and perhaps eventually lead to cell-based repair and neural regeneration.

## Acknowledgments

We thank C. Benod and P. Webb for assistance in early work; the late A. Sarrion-Perdigones for assistance with the dual luciferase reporter assay; R. Evans for sharing the *Tlx*^fl/fl^ mice; P. Yi for providing plasmids to express GST-NCOA3-RID; the BCM Center for Drug Discovery; the BCM NMR and Drug Metabolism Advanced Technology Core; M. Wang and J. Wang for access to GC-MS instrumentation, and C.-C. Lin and S.L. Holmes for technical support on GC-MS; J. Dannison and the MD Anderson Cancer Center Imaging Mass Spectrometry core for instruments and technical support. We thank Bert O’Malley, H. Zoghbi, M. Matzuk, P. Lucassen, and members of the Maletic-Savatic and Young laboratories for comments on the manuscript and helpful discussions; and V. Brandt for expert editing and comments on the work.

## Author Contributions

P.K. and F.S. contributed to experimental design, performed *in vitro* and *in vivo* experiments, respectively, analyzed the data, and drafted the manuscript. A.B. supplied human neural stem and progenitor cells and assisted in their analyses. D.B. and K.C. contributed to the IMS data acquisition and analyses. L.M. and K.R.M. performed the NMR experiments and analyses. A.C. and T.P. conducted luciferase and ALPHA screen assays. N.M. cloned the luciferase system. W.C. contributed first binding data. A.J-P., P.F-P., C.L.B. and E. E-P. contributed tools and data interpretation. S.C. synthesized fatty acid. I.O.R. conducted docking experiments. L.C. cloned and optimized the *in vitro* methods. K.V. supervised the luciferase experiments. D.D.M. contributed to the overall concepts, experimental design, and data interpretation. D.W.Y. and M.M-S. conceived and designed the experiments, oversaw the overall execution of the project, interpreted data, provided financial support, and wrote the manuscript. All authors discussed the results and commented on manuscript.

