## Supplemental Files for "Oleic acid triggers hippocampal neurogenesis by binding to TLX/NR2E1"

### Materials and Methods

**Human neural stem and progenitor cells (hNSPCs).** hNSPCs were generated from WA09 (H9) human embryonic stem cells (hESCs) maintained on matrigel-coated plates in Essential 8 Medium (E8) (*38*), using a variation of the dual SMAD inhibition protocol (*39-41*). First, hESCs were dissociated with Accutase. Two million cells were dispensed per well of an Aggrewell plate containing neural induction medium (NIM) and allowed to form aggregates. NIM was prepared from equal volumes of DMEM/F-12 and Neurobasal medium with 2% B27-supplement, 1% N2-supplement, 2 mM Glutamax, and 1% Pen/Strep. To promote cell survival, we used 10 µM Y-27632 (Tocris) during the initial 24 hr of culturing in Aggrewells (day 1). On day 2, we replaced three-quarters of the media and initiated SMAD inhibition by adding 10 µM SB-431542 (Tocris). Dorsomorphin (4 µM, Tocris) was then added to the culture from day 3 on. On day 5, we gently collected aggregates, sieved them through a reversible strainer, and transferred them into matrigel-coated plates with neural proliferation medium (NPM). NPM was prepared from equal volumes of DMEM/F-12 and Neurobasal media with 1% B27-supplement, 0.5% N2-supplement, 20 ng/mL bFGF, 20 ng/mL EGF, 2 mM Glutamax, and 1% Pen/Strep. To promote dorsalization, we added 10 µM cyclopamine (Tocris) to the media from day 6 on. Both SMAD inhibitors and cyclopamine were present in the media until day 9. We changed media daily until harvesting of rosette-shaped clusters of hNSPCs days 12-14, using Rosette Selection Reagent (*42*). After dislodging, rosettes were incubated in wells coated with 0.2% porcine gelatin to allow non-neural cells to differentially attach (*43*). We collected the floating fraction 1 hr following incubation, transferred them into non-coated cell culture flasks, and incubated them in suspension overnight. We plated rounded and floating hNSPCs spheres onto matrigel-coated 6-well plates the following day and propagated them until cells formed a confluent layer. All cultures were maintained in the presence of 1% Pen/Strep.

**hNSPC drug treatment and cell viability measurement.** One day before drug treatment, we plated 5,000 hNSPCs onto Matrigel coated 96-well plates (100µL/well). For fatty acid treatment, we prepared working stocks in DMSO and diluted them with hNSPC media before treating the cells. Following treatments, we performed MTT-based cell proliferation assay, adapted from ATCC. Briefly, we added 10 µL of MTT reagent (3-(4,5-dimethylthiazolyl-2)-2,5-diphenyltetrazolium bromide, 5mg/mL in phosphate-buffered saline (PBS)) into each well, followed by 100 µL of 10% SDS containing 0.01N HCl for cell lysis. After cell lysis, we measured absorbance from each well at 570 nm, using a TECAN plate reader. We used Prism 7 software from Graphpad Inc. (La Jolla, CA) for data analysis and normalization to the vehicle controls.
**NMR spectroscopy.** Proton NMR ($^1$H-NMR) was used to elucidate the structure of fatty acids abundant in hNSPCs. One million hNSPCs were collected at confluence, washed with PBS, and

spun down at 1,000×g for 1–2 min at 4°C to remove remnant culture media (*42*, *44*). Pellets were resuspended in 450 µL of PBS, pH 7.0 in $D_2O$ and 50 µL of 50 mM internal standard (2,2-dimethyl-2-silapentane-5-sulfonate (DSS) in $D_2O$. To ensure homogeneous cell suspension, samples were sonicated and kept at 4°C until data acquisition. Two-dimensional $^1H$-$^1H$ Correlation Spectroscopy (COSY) NMR was acquired for 24 hr at 25°C (800MHz NMR Avance, Bruker, Inc), using dipsi2esgpph (homonuclear Hartman-Hahn transfer) pulse sequence. Water suppression was achieved by excitation sculpting with gradients. Chemical shifts were referenced to DSS. The size of FID was k4k and k2k for F2 and F1 dimensions, respectively. The acquisition time was 0.18 sec for F2 and 0.09 sec for F1. The spectral width was 13.95 ppm. The line broadening for processing was 1.0 Hz. The number of scans was 24. All experiments were done in triplicate from the same culture (i.e., on the same day) to ensure reproducibility, with five biological replicates to ensure specificity. Special attention was paid to use powder-free gloves when handling NMR tubes to avoid fatty and other deposits from bare hands onto the glass that could potentially compromise data interpretation.

**Gas chromatography-mass spectrometry (GC-MS).** GC-MS was used to examine hNSPCs fatty acid content. hNSPCs were collected at confluence, washed with PBS, and spun down at 1,000 rpm for 3-5 min at 4°C to remove remnant culture media (*42*, *44*). Cell pellets were stored at -80°C until use. For fatty acid methyl ester (FAME) conversion, cell pellets were trans-esterified with 10% $BF_3$ in methanol at 75°C. The resulting methyl esters were extracted in hexane, dried under reduced pressure, re-dissolved in analytical LC-MS grade hexane, and analyzed by GC. GC analysis was carried out with a gas chromatograph equipped with a capillary column (Agilent 112-8867) and linked to an integrator. The column was temperature-programmed from 145 to 220°C at 2°C/min with an initial time of 26 min and a final time of 1 min. Helium carrier gas and a split ratio of 100:1 was used. Fatty acid peaks were identified by comparison with authenticated standards (Nu-check, catalogue#566C).

**Imaging Mass Spectrometry (IMS).** Imaging mass spectrometry enables untargeted visualization of the spatial distribution of metabolites in tissue section by extracting an individual mass-to-charge (*m*/*z*) value from each pixel's spectrum. The metabolite identity is discerned by matching of its mass to a database of known molecules within a certain mass-error range. To map lipids in the mouse brain, we euthanized a three-month-old wild-type C57BL/6J mouse with isoflurane overdose, dissected its brain, and placed it into a Tissue-Tek® paraffin-embedding cassette (Electron Microscopy Sciences, Hatfield, PA) pre-cooled in liquid nitrogen for 30 sec. Tissue was first held ~5 cm above liquid nitrogen level for 45 sec and then slowly dipped into liquid nitrogen for 5 sec followed by retraction for 5 sec. This was repeated two more times, and the brain was then wrapped in pre-cooled foil (in liquid nitrogen) and placed into pre-cooled glass bottles at -80°C until use[40]. Tissue was mounted onto a cryostat with a drop of $ddH_2O$ and cryosectioned into 10 µm slices. Slides were kept at -80°C until data acquisition. Matrix Assisted Laser Desorption/Ionization (MALDI)-IMS was performed using Waters Synapt G2-Si High Definition Mass Spectrometer equipped with a MALDI source for IMS. The slide with the brain section was first thawed to room temperature under vacuum, ensuring that no water droplets formed on the surface (*42*). The tissue section was coated with 9-aminoacridine (9AA; 600 mg) as the MALDI matrix for 20 min, using an automated sublimation matrix applicator (Shimadzu IM Layer). The coated slides were rehydrated in a heated humidifying chamber for 3 min, using 200 µL of 10% methanol.

Prior to acquiring the data, the instrument was calibrated for mass accuracy using red phosphorus, calibrating the mass range from m/z 50-2000. Before loading into the mass spectrometer, the slide with tissue section was scanned using an EPSON scanner for mapping areas of interest into the High Definition Imaging (HDI, Waters Corporation) 1.4 software. The laser power was set to 250 (arbitrary units) with 300 laser shots per pixel data. The laser raster step was set to 60 µm to match the laser spot size and the ionization source was set to negative ionization mode. Run time was about 130 min. After acquisition, the raw file was imported into HDI 1.4 software, where the data were processed into a collection of images (*45*). After a table of mass lists was generated from the data, we selected a particular mass identifying a metabolite of interest to display a heatmap distribution of the signal. To determine signal intensities in particular regions, we selected regions of interest over the generated images. The signal intensities were normalized by total ion count (TIC) to account for varying ionization efficiencies across the tissue sample. The m/z values of different fatty acids of interest were compiled using the Human Metabolome Database (HMDB).

**Modeling the TLX ligand-binding domain and ligand docking.** We modeled the TLX ligand binding domain (TLX LBD) (PDB code: 4AXJ) after fatty acid ligand-bound HNF4α (PDB code: 1M7W) using SWISS-MODEL Workspace. Docking was performed using Autodock Vina run through PyRx to manage the workflow. Docking outputs were visualized using PyMoL. 18:1ω9 structure was prepared by generating an energy-minimized 3D structure in ChemBioDraw3D. This was followed by processing with Autodock Tools 1.5.4 using the "make ligand" function. TLX LBD homology model was also processed with Autodock Tools 1.5.4 using the "make macromolecule" function. Docking runs were performed within a 25–30 Å cubic search space surrounding the binding pocket. Outputs were selected according to their ranking in PyRx and how their poses matched that of the ligand occupying the binding pocket of HNF4α.

**Cloning, expression, and purification of TLX LBD, GST-tagged NCOA3 (SRC3) receptor interaction domain (RID), and coregulator peptide synthesis.** We cloned human TLX LBD (residues 181-385) into pMCSG7 and pMCSG51 protein expression vectors and verified by DNA sequencing. 6xHis-TLX LBD was expressed from pMCSG7 vector in *E. coli* by overnight IPTG induction at 16-18°C. The cell pellet was frozen and lysed before protein purification by sonication. To obtain biotinylated TLX LBD, 6xHis-AviTag-TLX LBD was expressed from pMCSG51 vector in the presence of 50 µM biotin. 6xHis-TLX LBD and 6xHis-AviTag-TLX LBD were purified by affinity chromatography, dialyzed to remove imidazole, and further purified by size exclusion chromatography as reported (*17*), using BioRad NGC protein purification system. GST-NOCA3-RID was expressed and purified as described (*46*), using BioRad NGC protein purification system. Corepressor atrophin peptide with conserved **AL**XX**L**XX**Y** Atrobox containing motif (amino acid sequence PPYADTP**AL**RQ**L**SE**Y**ARPHVAFSP, ≥95% purity) was custom-synthesized with and without N-terminal biotin-tag (GenScript, USA). Similarly, nuclear receptor coactivator1 peptide (NCOA1-II peptide) with consensus canonical **L**XX**LL** coactivator motif (amino acid sequence: LTERHKI**L**HR**LL**QEGSPSD, ≥95% purity) was custom-synthesized with and without N-terminal biotin-tag (GenScript, USA).

**Biolayer interferometry assays (BLI).** We used BLI, an optical label-free technology, to examine interactions between the TLX LBD and different fatty acids. BLI binding measurements were performed with 6xHis-TLX LBD using the Octet Red 96 instrument (Ni-NTA Biosensors, catalogue #18-5103, *FortéBio Inc./Molecuar Devices LLC*, USA). Dip and read Ni-NTA biosensors prepped with assay buffer (150 mM NaCl, pH 7.0, 1 mM DTT and 2 mM CHAPS) and TLX LBD were immobilized on the sensors to saturation. The sensors were equilibrated to appropriate vehicle control prepared in buffer for initial reading before measuring the binding response at increasing concentrations of various analytes diluted in the buffer. Data analysis was performed using *ForteBio* software to obtain the equilibrium response and plotted using Prism 7 software from Graphpad Inc. (La Jolla, CA) to obtain the corresponding equilibrium dissociation constant ($K_D$) under these experimental conditions.

**ALPHA screen assay.** The ALPHA screen measures the proximity of donor and acceptor beads using luminescence: if the two beads are close, a singlet oxygen is transferred from the donor to the acceptor bead, inducing an increase in luminescence. Assays were performed following the manufacturer's protocol (AlphaScreen Histidine (Nickel Chelate) and AlphaScreen GST Detection Kit, catalogue# 6760619C & 6760603C, PerkinElmer, USA) with some modifications. Fatty acids were tested for TLX-atrophin peptide binding in 20 mM Tris, 300 mM NaCl, pH 7.5, 1 mM DTT or TCEP and 0.03% CHAPS. Stock fatty acids in 100% DMSO were dissolved and diluted in assay buffer and 5 µL was added to the assay plate. Biotinylated atrophin peptide (200 nM final concentration) and 6xHis-TLX LBD (1000 nM final concentration) were mixed and incubated for 10 min. Donor and acceptor beads (10 µg/mL final concentration) were added to the protein-peptide mix, incubated for 10-15 min each, and 15 µL of the master mix was added to each well under low light for ligand screen.

We performed a competition experiment with atrophin peptide without biotin tag under similar conditions, to measure the half-maximal effective concentration ($EC_{50}$) for the corepressor binding without any ligands. The plates were spun down, mixed with a shaker, incubated for 60 min, and read on a TECAN plate reader using the manufacturer's protocol. To examine the interaction of fatty acids with the coactivator, assays were performed in 50 mM sodium phosphate, 150 mM NaCl, pH 7.0, 1 mM DTT or TCEP and 0.03% CHAPS. Biotinylated SRC peptide (100 nM final concentration) and 6xHis-TLX LBD (500 nM final concentration) were mixed with the donor and acceptor beads (10 µg/mL final concentration) and 15 µL of the mix was added to the plate with 5µL of fatty acids. The mixture was incubated for 2 hr before reading the signal. We performed a competition experiment with NCOA1-II peptide without biotin tag under similar conditions, to obtain $EC_{50}$ for the NCOA1-II binding in the presence of 200 µM oleic acid, and a similar competition experiment with NCOA1-II peptide without biotin tag was performed to measure $EC_{50}$ for the GST-NCOA3-RID binding to 6xHis-AviTag-TLX LBD in the presence of 200 µM oleic acid. $EC_{50}$ was calculated by plotting the DMSO-normalized ALPHA signal (nonlinear regression fit), using Prism 7 software.

**Homogenous Time-Resolved Fluorescence (HTRF) assay.** HTRF is a resonance energy transfer in which long emission fluorophores (lanthanides) are used as donors. The comparison measurement of the two emitted wavelengths over time is calculated for a HTRF response. The HTRF assay was carried out in low-volume 384-well plates at room temperature using Anti-His-Terbium (Tb) antibody and streptavidin-d2 (CisBio/PerkinElmer, USA). Fatty acids or DMSO were prepared in assay buffer (50 mM sodium phosphate, 150 mM NaCl, 1 mM DTT, 100 µM CHAPS, 10% Glycerol, pH 7.0) and 2.5 µL was added to the assay plate. A 7.5 µL mixture of 6xHis-TLX LBD (10 nM final concentration), Tb donor (1x final), biotinylated SRC1-II peptide (100 nM final concentration) and d2 acceptor (25 nM final concentration) was added to 384-well plate. Plates were sealed and incubated at room temperature for 30 min. The Tb donor was excited at 340 nm, its emission was monitored at 665 nm, and the d2 acceptor emission was measured at

620 nm. To determine $EC_{50,}$ data were normalized to vehicle control and fitted using nonlinear regression fit using Prism 7 software.

**Cell-based dual-luciferase reporter assay.** We cloned TLX-based luciferase reporter plasmids using Goldenbraid 2.0 (GB2.0), a multipartite, single-pot assembly cloning method utilizing the type IIs restriction enzymes *Bsa*I and *Bsm*BI for iterative synthetic assembly (*47, 48*), adapted to luciferase assay (*49*). We generated a vector expressing TLX isoform b, pCMV-TLXb, from DNA parts encoding the cytomegalovirus (CMV) enhancer-promoter, TLX coding DNA sequence (CDS), and a poly(A) termination signal from the bovine growth hormone gene (bGH). Briefly, cloning involved mixing 40 ng of GB2.0-compatible DNA part vectors for the CMV promoter (Addgene #118048), TLX CDS, and bGH terminator (Addgene #118061) with 75 ng of destination vector pColE1_Alpha1 (Addgene #118044), 1 µl of *Bsa*I enzyme (New England Biolabs) and 1 µl of T4 DNA Ligase (Promega), and 2 µl of ligase buffer (Promega) in a single tube. Assembly was performed using a standard thermocycler with cycling conditions of 2' at 37 °C and 5' at 16 °C steps for 25-50 cycles (*47*).

The luciferase reporter plasmid, p3xTAE_FLuc_Renilla_sfEGFP, consisted of three transcription units cloned into the destination vector pColE1_Alpha2 (Addgene #118045). Each unit was cloned using GB2.0 cloning similarly to pCMV_TLX cloning as above. The three transcriptional units (TUs) were: (1) a TLX responsive firefly luciferase (FLuc) CDS, (2) constitutively expressed Renilla luciferase, and (3) constitutively expressed superfolder green fluorescent protein sfEGFP (*50*). The firefly luciferase TU, p3xTAE_FLuc, consisted of a synthetic TLX-responsive enhancer element made of a triplicate of TLX-activating element (TAE) (*31*), upstream of a synthetic minimal promoter, MiniP (Addgene #118049), FLuc CDS (Addgene #68201) and bGH terminator assembled into pColE1_Alpha2. The Renilla TU, pSV40_Renilla, was made of the viral SV40 promoter, Renilla luciferase CDS (Addgene # 118060) and bGH terminator (Addgene #118061) cloned into pColE1_Alpha1. Finally, the sfEGFP TU, pSV40_sfEGFP, consisted of the SV40 promoter, sfEGFP, and bGH terminator (Addgene #118061) cloned in pColE1_Alpha2. Assembly of p3xTAE_FLuc_Renilla_sfEGFP involved cloning $p(A)^n$-PAUSE, a transcriptional insulator (Addgene #118069) upstream of p3xTAE_Fluc into the destination vector pColE1_Omega1 (Addgene #118046) and assembly of pSV40_Renilla and pSV40_sfEGFP into pColE1_Omega2 (Addgene #118047)  GB2.0 reactions were performed as above in separate *Bsm*BI (New England Biolabs) assembly reactions to produce intermediates pPAUSE-3xTAE_FLuc and pSV40_Renilla_sfGFP. These intermediates were then further assembled together to produce the final product in a *Bsa*I driven assembly reaction as above.

Molecular biology experiments, including plasmid maps and *in silico* experimentation, were designed and generated using SnapGene software (GSL Biotech LLC) (http://www.snapgene.com/products/snapgene/). Standard microbiology techniques were used for chemical transformation of the *E. coli* strain K12/DH10B cells (Thermo Scientific). DNA was recovered using QIAprep Spin Miniprep Kit (QIAGEN). Assembly products were confirmed via restriction enzyme DNA fingerprinting and visualized using agarose gel electrophoresis. HeLa cells were seeded in 5% charcoal stripped media and incubated at 37°C for 24 hrs.

To test the effects of different fatty acids on TLX using the dual luciferase reporter, we used HeLa cells obtained from the tissue culture core at Baylor College of Medicine. They were passaged and maintained in Dulbecco's Modified Eagle Medium (DMEM) cell culture media supplemented with 10% fetal bovine serum (FBS) and 1% Pen-Strep, following American Type Culture Collection (ATCC) guidelines. For luciferase experiments, we plated 3 million HeLa cells into 6-well cell culture flasks in 5% stripped FBS DMEM media without phenol red. After 24 hr, cells were transfected using lipofectamine reagent (ThermoFisher, USA, catalogue # L3000008) and 1:1 TLX reporter plasmid (2 µg):control, or TLX expression plasmid (2 µg). 24 hr later, cells were trypsinized and plated to 24-well plates, and fatty acids were added to 1% charcoal-stripped FBS media. We lysed cells 24 hrs following fatty acid treatment using Promega 1x-passive lysis buffer. To examine luminescence, cell lysate was transferred to a 384-well white plate after addition of firefly and renilla substrates (Promega, US, Dual-Luciferase® Reporter Assay System, catalogue # E1980), using the CLARIOstar microplate reader. We normalized firefly luciferase to renilla luciferase signal from each well and normalized response was plotted using Prism 7 software.

Similarly, we tested different fatty acids with TLX after nucleofection into HeLa cells with the dual luciferase reporter system. Following transfection program I-013 (Amaxa Nucleofector Technology, Lonza, US), 5 million HeLa cells were electroporated with 2 µg of TLX reporter plasmid and 2 µg of TLX expression plasmids and grown in a culture flask with 5% stripped FBS DMEM media without phenol red. After 24 hr, cells were trypsinized and plated into 24 well-plates, and fatty acids were added to 1% stripped FBS DMEM media without phenol red. After 24 hrs of fatty acid treatment, the cells were lysed and luciferase response was measured as above.

**Transgenic mice.** *Lfng*-eGFP mice (RRID:MMRRC_015881-UCD) were obtained from GENSAT (*51*) and received as FVB/N-C57BL/6 hybrids. They were crossed to C57BL/6J mice for at least 10 generations and fully characterized prior to use (*32*). *Tlx*$^{loxp/loxp}$ mice were a gift from Dr. Ronald Evans (*13*). C57BL/6J mice and the AI14 (RCL-tdT) reporter line were from The Jackson Laboratory (Bar Harbor, ME) (JAX 000664; RRID:IMSR_JAX:000664 and JAX 007908; RRID:IMSR_JAX:007908, respectively) (*52*). *Lfng*-CreER$^{T2}$ mice were generated in the Maletic-Savatic laboratory (*32*). All experiments were done according to the Baylor College of Medicine Institutional Animal Care and Use Committee approved protocol (AN-5004).

Tamoxifen (120 mg in 10 mL of 1:9 ethanol:corn oil mixture) solution was administered intraperitoneally to *Lfng*-CreER$^{T2}$-based mice. Control mice were injected with ethanol:corn oil mixture only. SCD inhibitor (CAY10566) (Cayman Chemical, Ann Arbor MI, cat#10012562; 3 mg/kg body weight) dissolved in 0.03 N HCl was given orally daily for 5 consecutive days followed by 4X EdU (2 hr apart) on the day of sacrifice. BrdU (150 mg/kg) was administered intraperitoneally. CldU (85 mg/kg) and IdU (115 mg/kg) were administered in equimolar concentrations. BrdU and CldU were dissolved in sterile saline. IdU was dissolved in sterile saline solution that contained 2% of 0.2 N NaOH. EdU (50 mg/kg) was dissolved in sterile saline solution and detected with Click-iT™ EdU Alexa Fluor™ 647 Imaging Kit (Thermo Fisher Scientific).

**Stereotactic injections.** Mice were anesthetized with Rodent III combo (37.5 mg/mL ketamine, 1.9 mg/mL xylazine and 0.37 mg/mL acepromazine) and received a single dose of analgesic Buprenorphine Sustained Release (5mg/mL) (Zoopharm, Windsor, CO) subcutaneously. After positioning the mice within the stereotactic apparatus, we delivered fatty acids into the right dentate gyrus at three sites relative to Bregma: Anteroposterior (AP) -1.5 mm, -1.7 mm, or -1.9 mm, Lateral-Lateral (LL) -1.6 mm, and Dorsoventral (DV) -1.9 mm, using nanoinjector (Nanoject II, Drummond Scientific, Broomal, PA). Each site received 305.4 nL of fatty acids in 6 pulses (50.9 nL per pulse). Injection was done in a slow delivery mode over 2 sec per pulse. Each pulse was separated by 15 sec. Glass capillary was gradually moved in and retracted from the tissue, over 3 min. The same coordinates were used for sham injections into the left dentate gyrus. Post-operative care was done as per IACUC-approved protocol.

**Immunohistochemistry.** Mice were perfused transcardially with 30 mL of PBS followed by 30 mL of 4% (w/v) ice cold paraformaldehyde (PFA) in PBS. Brains were post-fixed in 4% PFA solution overnight at 4°C. PFA was then replaced with PBS and tissues were kept at 4°C for further use. Free-floating serial 50 µm sagittal sections were cut using Vibratome 1500 and collected in five parallel sets, each containing 14 sections 250 µm apart from each other. Sections were immunostained as described (*32*).

For immunostaining against BrdU, CldU, and IdU, sections were treated with 2N HCl for 30 min at 37°C, followed by rinsing with PBS and incubation with 0.1 M sodium tetraborate (pH 8.5) for 10 min at room temperature and then again rinsing with PBS. For other antigens, sections were incubated with primary antibodies overnight at 4°C after initial permeabilization and blocking at room temperature for 2 hrs. Sections were then washed three times with PBS and incubated with fluorochrome-conjugated secondary antibodies for 2 hrs at room temperature. Sections were washed three times with PBS and mounted on slides with DakoCytomation Fluorescent Mounting Medium (DakoCyomation, Carpinteria, CA).

EdU staining was performed according to the manufacturer's protocol (Click-iT EdU-Alexa-Fluor$^{TM}$ 647 Imaging Kit, ThermoFisher, Waltham, MA). The eGFP signal from *Lfng*-eGFP and tdTomato signal in Ai14-crossed control and conditional knockout mice was amplified with antibodies against GFP (chicken anti-GFP; Aves Labs Cat# GFP-1020 RRID:AB_10000240, at 1:1000) or against RFP rabbit anti-RFP (Rockland Cat# 600-401-379 RRID:AB_2209751, at 1:500); goat anti-RFP (SICGEN, Cantanhede, PORTUGAL, at 1:200), respectively, following antigen retrieval with HCl treatment. For other antigens, the following antibodies were used: rabbit anti-Dcx (Cell Signaling Technology Cat# 4604S RRID:AB_10693771) at 1:200; mouse anti-GFAP (Sigma-Aldrich Cat# G3893 RRID:AB_477010) at 1:1000; rabbit anti-GFAP (Dako Cat# Z0334 RRID:AB_10013382) at 1:1000, mouse anti-NeuN (Millipore Cat# MAB377 RRID:AB_2298772) at 1:500, rabbit anti-Sox2 (Abcam Cat# ab97959 RRID:AB_2341193) at 1:500; mouse anti-BrdU (Bio-Rad / AbD Serotec Cat# OBT0030CX RRID:AB_609566) (used for detecting CldU) at 1:300; rat anti-BrdU (Becton Dickinson and Company Cat# 347580 RRID:AB_10015219) (used to detect IdU); secondary antibodies (conjugated with Alexa 488, 594, or 657) (all pre-absorbed against other species to prevent cross reactivity) (Jackson Immunoresearch, West Grove, PA) at 1:500. Sections were counterstained with DAPI (5 µg/mL, Sigma) at 1:1000.

20 µm optical sections were scanned with confocal microscopy (Zeiss LSM 710). 3D reconstruction and orthogonal views were acquired via ZEN2012 SP1 software (Zeiss, Thornwood, NY). Cells in the uppermost focal plane of the dentate gyrus were excluded from quantification. Total counts from 14 sections were multiplied by 10 to get the total number of cells in the two dentate gyri. We used the optical dissector method to quantify and characterize proliferating cell types (*32*, *53*). NSCs were identified based on *Lfng*-eGFP$^+$ and/or *Lfng*-CreER$^{T2}$;RCL-tdT$^+$ triangular Sox2$^+$ nuclei located in the subgranular zone (SGZ) with a GFAP$^+$ radial process spanning the granule cell layer and ending with fine eGFP$^+$ arborizations in the granule cell layer/molecular layer boundary (*54*). In C57BL/6J wild-type mice, NSCs were identified as cells with the triangular soma in the SGZ and a GFAP$^+$ radial process originating from a Sox2$^+$ nuclei in SGZ and spanning throughout granule cell layer. ANPs were identified as GFAP$^-$ Sox2$^+$ round cells located in the SGZ. Neuroblasts and immature neurons were identified as Dcx$^+$ cells with single or multiple processes. Granule neurons were identified as NeuN$^+$. Ratio of EdU$^+$, CldU$^+$, or IdU$^+$ cells among a certain cell type (NSC, ANP etc.) was calculated by dividing the EdU$^+$, CldU$^+$, or IdU$^+$ cells to the total number of the respective cell type.

**Fluorescence Activated Cell Sorting (FACS).** Mice were euthanized with isoflurane and perfused transcardially with 10 mL of ice-cold PBS. Brains were immediately transferred into a culture dish containing ice-cold HBSS. Dentate gyri were isolated as described (*55*) and placed into 2mL of Hybernate EB (HEB) complete media (BrainBits, Springfield, IL) and 2mL of papain solution (heat-activated at 37°C) for 2 min at 37°C. Samples were passed through 1 mL pipette tips 2-3 times to break up the tissue and further incubated at 37°C for 18 min, gently swirling every 5-6 min to ensure enzyme access to the whole tissue. Papain solution was replaced with HEB media and tissue was gently triturated through the fire polished and salinized Pasteur pipette for 10-15 passes or until about 85% of tissue dissociation was achieved. After allowing the tissue debris to settle for 1 min, we transferred supernatant containing dispersed cells to a new tube after passing through a 70 μm cell strainer (Corning, Durham, NC). Cells were centrifuged at 200 rcf for 3 min and the supernatant containing the debris was discarded. Cells were re-suspended in 0.5 mL of pre-warmed low fluorescence Hibernate E media (BrainBits, Springfield, IL) by gently pipetting 20 times and once more passed through the 70 μm cell strainers. Sytox$^{TM}$ Red (ThermoFisher, Waltham, MA) dead cell stain was added to discard non-viable cells during FACS.

For FACS calibration, all control tubes (tdTomato+, eGFP+, SytoxTM Red+, wild type) were prepared in advance. A 70 μm nozzle at 70 psi was used for the sorting (BD FACS Aria II). All tubes were kept chilled and protected from light until FACS procedure. tdTomato+ and eGFP+ (TLX mutant NSC clones from *Tlx*$^{fl/fl}$ mice), only tdTomato+ (TLX mutant NSC progeny), only eGFP+ (wild-type NSC clones from *Tlx*$^{fl/fl}$ mice), tdTomato- and eGFP- (wild-type non-NSCs) cells from 18:1ω9 injected *Tlx*$^{fl/fl}$/*Lfng*-eGFP/*Lfng*-CreER$^{T2}$/RCL-tdT mice and eGFP+ (NSCs) or eGFP- (non-NSCs) cells from 18:1ω9 or 18:3ω3 injected *Lfng*-eGFP mice were sorted. Each sample was generated from 2 dentate gyri. Sorted cells were kept chilled in RNAprotect cell reagent (Qiagen, Germantown, MD) during FACS procedure and at -20°C until RNA isolation.

**RNA isolation, whole transcriptome amplification, and RT-PCR.** RNA was isolated from sorted cells using RNeasyplus Micro kit (Qiagen, Germantown, MD). To eliminate possible DNA contamination, we used RNase free DNase set (Qiagen, Germantown, MD). We used QuantiTect Whole Transcriptome Kit (Qiagen, Germantown, MD) for reverse transcription and preamplification of isolated RNA. RT-PCR was performed using RT2 profiler PCR array Mouse Neurogenesis (Qiagen, cat# PAMM-404Z) Mouse Cell Cycle (Qiagen, cat# PAMM-020Z), Mouse Notch Signaling Pathway (Qiagen, cat# PAMM-059Z) and RT2 SYBR Green qPCR master mix (Qiagen, cat# 330513). Expressions were normalized to β-2 macroglobulin (B2m) gene. We calculated fold changes compared to control group using the ΔΔCt method. Genes upregulated (ΔΔCt ≥2 or ≥4-fold change) in 18:1ω9-treated or 18:3ω3-treated mice in comparison to sham-injected wild type mice were identified. To identify genes suppressed by TLX, we compared sham-injected *Lfng*-eGFP+ NSCs to sham-injected *Tlx* mutant NSC clones (eGFP+; tdTom+) in *Tlx*$^{fl/fl}$ mice. Genes upregulated in the mutant clones were accepted as genes suppressed by Tlx. To identify genes regulated by 18:1ω9 in a TLX-independent manner, we compared 18:1ω9-treated *Tlx* mutant NSC expression to sham-injected *Tlx* mutant NSCs.

**Statistical analysis.** Statistical analysis was performed using GraphPad Prism 8.0 (GraphPad RRID:SCR_002798). The sample size was determined based on published data (*56*) (*57*). Experiments involving 2 groups were compared using un-paired Student t-test. Experiments involving more than 2 groups with one variable were compared by One-Way Analysis of Variance (ANOVA), or Two-way ANOVA followed by Tukey HSD post-hoc test analysis for pairwise comparisons. Significance was defined as $p<0.05$.

**Methods for chemical synthesis.** Fatty acids were purchased from commercial sources except for 18:1ω5 and *trans*18:1ω5, which were synthesized in-house. All starting materials and chemical reagents were purchased from commercial sources and used without further purification for the synthesis of 18:1ω5 and *trans*18:1ω5. Solvents were purchased as either anhydrous grade products in sealed containers or reagent grade and used as received. All reactions were carried out in dry glassware under a nitrogen atmosphere using standard disposable or gastight syringes, disposable or stainless-steel needles, and septa. Stirring was achieved with magnetic stir bars. Flash column chromatography was performed with $SiO_2$ (230-400 mesh) or by using an automated chromatography instrument with an appropriately sized column. Thin layer chromatography was performed on silica gel $60F_{254}$ plates (E. Merck). Non-UV active compounds were visualized on TLC using one of the following stains: $KMnO_4$, bromocresol, *p*-anisaldehyde. $^1$H and $^{13}$C NMR spectra were recorded on an instrument operating at either 600 MHz, or 151 MHz, respectively. All NMR chemical shifts are quoted on the δ scale and were referenced to residual non-deuterated solvent as an internal standard. Signal multiplicities are described using the following abbreviations: s=singlet, d=doublet, t=triplet, b=broad, quar=quartet, quin=quintet, m=multiplet, v=very; abbreviations are combined, *e.g.* vbs=very broad singlet.

**Chemical synthesis of 18:1ω5.** Chemical synthesis of cis and trans 18:1ω5 fatty acids were achieved as schematically shown below by adopting literature protocols.

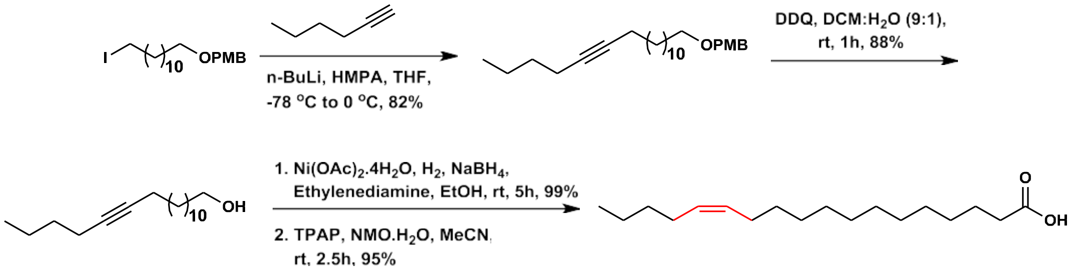


**Synthesis of alkyne intermediate.** The alkyne intermediate was synthesized by adopting a previously known method (*58*). Into an oven dried round bottom flask equipped with magnetic stir bar and septum, 1-hexyne (1 equiv.) was dissolved in dry THF (5 mL/mmol) and cooled to -78°C. Next, n-BuLi (1.5 equiv., 1.6 M solution in hexanes) was added and the mixture was stirred for 25 min followed by the addition of 1-(((12-iodododecyl)oxy)methyl)-4-methoxybenzene (2 equiv.). The reaction was allowed to warm to 0 °C over 1.5 hr and then warmed to ambient temperature and stirred for an additional 1 hr. Water (20 mL) was added to the reaction mixture followed by $Et_2O$ (50 mL). The layers were separated, and the aqueous layer was washed with $Et_2O$ (50 mL). The combined organic layers were dried ($Na_2SO_4$) and concentrated. The crude residue was purified by silica gel chromatography (EtOAc/hexanes, 5:95) to provide the pure product.

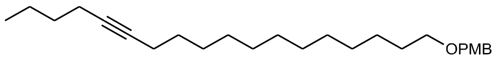


**1-methoxy-4-((octadec-13-yn-1-yloxy)methyl)benzene;** Molecular Formula: $C_{26}H_{42}O_2$; Rf (10% ethyl acetate/hexanes): 0.3; **$^1$H NMR (600 MHz, Chloroform-*d*)** δ 7.30 – 7.27 (m, 2H), 6.90 (d, $J$ = 8.6 Hz, 2H), 4.46 (s, 2H), 3.83 (s, 3H), 3.46 (t, $J$ = 6.7 Hz, 2H), 2.20 – 2.13 (m, 4H), 1.65-1.58 (m, 2H), 1.55 – 1.45 (m, 4H), 1.46 – 1.40 (m, 2H), 1.39–1.34 (m, 4H), 1.33–1.25 (m,

12H), 0.93 (t, *J* = 7.2 Hz, 3H); **$^{13}$C NMR (151 MHz, $CDCl_3$)** δ 159.10, 130.85, 129.19, 113.75, 80.22, 80.16, 72.50, 70.25, 55.26, 31.29, 29.79, 29.60, 29.55, 29.17, 28.87, 26.22, 21.93, 18.77, 13.63.

**PMB Deprotection.** The p-methoxybenzyl (PMB) protecting group was removed from the alkyne to obtain the free alcohol intermediate by following a previously known method (*59*). Into a round bottom flask equipped with magnetic stir bar and septum, the alkynyl ether compound (1 equiv.) was dissolved in dichloromethane: water (10 mL:1 mL per mmol). Next, DDQ (4 equiv.) was added and the reaction was allowed to stir at room temperature for 1 hr, after which time the TLC showed complete consumption of the starting material. The mixture was diluted with DCM and washed with saturated aqueous $NaHCO_3$ solution. The organic phase was collected and dried over anhydrous Na2SO4 and the solvent was removed under reduced pressure to give the crude residue. Purification by silica gel chromatography (EtOAc/hexanes, 10:90) provided the pure product.

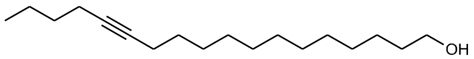


**octadec-13-yn-1-ol;** Molecular Formula: $C_{18}H_{34}O$; Rf (20% ethyl acetate/hexanes): 0.2; **$^1$H NMR (600 MHz, $CDCl_3$)** δ 3.64 (t, *J* = 6.7 Hz, 2H), 2.17–2.11 (m, 4H), 1.60 – 1.54 (m, 2H), 1.46 (m, *J* = 11.2, 7.0, 4.1, 3.6 Hz, 4H), 1.42 – 1.32 (m, 6H), 1.3–1.25 (m, 12H), 0.90 (t, *J* = 7.2 Hz, 3H); **$^{13}$C NMR (151 MHz, $CDCl_3$)** δ 99.98, 80.22, 63.11, 32.83, 29.61, 29.59, 29.54, 29.44, 29.18, 29.16, 25.74, 21.93, 18.76, 18.45, 13.63.

**Reduction of alkyne to cis alkene.** The deprotected alkyne was reduced to cis alkene by adopting a previously known method (*60*). Into a round bottom flask equipped with magnetic stir bar and septum, $Ni(OAc)_2$•$4H_2O$ (0.32 equiv.) was suspended in 2 mL of ethanol under a $H_2$ atmosphere and cooled to 15 °C and NaBH4 (0.78 equiv.) was added as an ethanolic solution (1 M solution ). After 10 minutes, ethylenediamine was added (3.55 equiv.) in a solution of ethanol (0.5 mL) and the mixture was stirred an additional 10 minutes before adding the alkyne (1 equiv.) as a solution in ethanol (1 mL). Before and after each addition, three cycles of vacuum/$H_2$ were applied. The reaction was stirred under $H_2$ atmosphere for 5 hr. The mixture was then filtered through Celite® and rinsed with EtOAc. The solvent was removed under reduced pressure and the crude residue was purified by silica gel chromatography (EtOAc/hexanes, 10:90) to provide the pure product.

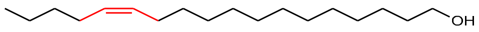


**(Z)-octadec-13-en-1-ol;** Molecular Formula: $C_{18}H_{36}O$; Rf (25% ethyl acetate/hexanes): 0.2; **$^1$H NMR (600 MHz, $CDCl_3$)** δ 5.40–5.34 (m, 2H), 3.65 (t, *J* = 6.7 Hz, 2H), 2.05–2.01 (m, 4H), 1.61 – 1.54 (m, 2H), 1.53 (s, 1H), 1.36–1.32 (m, 8H), 1.30–1.26 (m, 11H), 0.91 (t, *J* = 7.3 Hz, 3H); **$^{13}$C NMR (151 MHz, $CDCl_3$)** δ 129.88, 129.82, 63.03, 32.81, 31.97, 29.77, 29.63, 29.60, 29.55, 29.31, 27.19, 26.91, 25.75, 22.34, 13.97.

**Alcohol oxidation to carboxylic acid.** 18:1ω5 fatty acid was obtained from the conversion of the alcohol intermediate to the carboxylic acid by adopting a previously known (*61*) method.
Into a round bottom flask equipped with magnetic stir bar and septum, the alcohol compound (1

equiv.) was dissolved in acetonitrile (5 mL/mmol) followed by the addition of N-methyl morpholine N-oxide (NMO) monohydrate (10 equiv.) and tetrapropylammonium perruthenate (TPAP) (10 mol%). The reaction was allowed to stir at room temperature for 2.5 hr, after which time TLC indicated complete consumption of starting material. The reaction was quenched by the addition of excess of 2-propanol and the volatiles were evaporated under reduced pressure. The residue was redissolved in EtOAc (5 mL) and the mixture was filtered through Celite® and rinsed with EtOAc. The solvent was removed under reduced pressure and the crude residue was purified by silica gel chromatography (EtOAc/hexanes, 50:50) to provide the pure product.

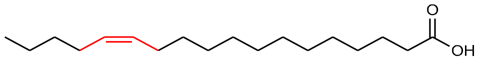


**(Z)-octadec-13-enoic acid;** Molecular Formula: $C_{18}H_{34}O_2$; Rf (50% ethyl acetate/hexanes): 0.2; **$^1$H NMR (600 MHz, $CDCl_3$)** δ 5.37–5.32 (m, 2H), 2.35 (t, *J* = 7.5 Hz, 2H), 2.05–1.98 (m, 4H), 1.64–1.61 (m, 2H), 1.39 – 1.22 (m, 20H), 0.92 – 0.86 (m, 3H); **$^{13}$C NMR (151 MHz, $CDCl_3$)** δ 180.31, 129.89, 129.84, 34.10, 31.98, 29.60, 29.58, 29.54, 29.44, 29.31, 29.24, 29.07, 27.20, 26.92, 24.68, 22.35, 13.99.

**Chemical synthesis of *trans*18:1ω5.**

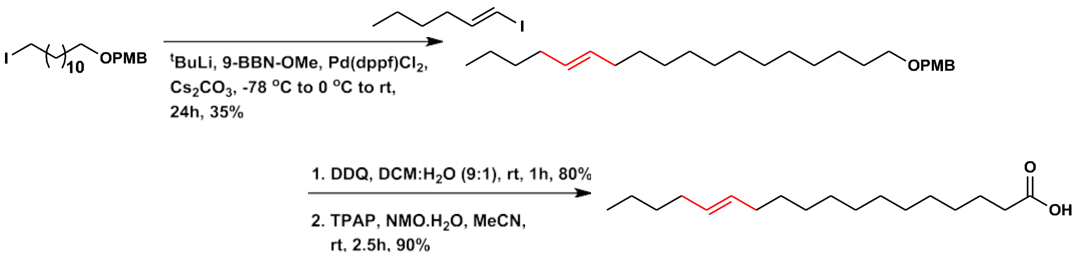


**Synthesis of trans alkene.** The PMB protected trans-octadec-13-en-1-ol was synthesized by adopting a previously known method (*62*). Into a round bottom flask equipped with magnetic stir bar and septum under nitrogen, the alkyl iodide (1 equiv.) was dissolved in $Et_2O$ (5 mL/mmol) and cooled to -78 °C. Next, t-BuLi (1.9 M in pentane, 2.5 equiv.) was added dropwise via syringe, followed by MeO-9-BBN (1 M in hexanes, 2.5 equiv.) and THF (10 mL/mmol). The resulting cloudy mixture was warmed to room temperature and stirred for 1 hr. Next, 1.27 mL of a 3.0 M solution of $Cs_2CO_3$ in $H_2O$ was added followed by the vinyl iodide (0.8 equiv.) as a solution in DMF (1 mL) and $Pd(dppf)Cl_2$ (5 mol%). The resulting red/brown suspension was covered with aluminum foil, and the reaction was stirred at room temperature for 20 hr. The reaction mixture

was diluted with $H_2O$ (10 mL) and ether (50 mL), the layers were separated, and the aqueous layer was extracted with ether (30 mL). The combined organic layers were washed with brine (1 x 50 mL), dried over $Na_2SO_4$, and concentrated under reduced pressure to give the crude residue. Purification by silica gel chromatography (EtOAc/hexanes, 5:95) provided the pure product. PMB deprotection was performed according followed by oxidation of trans-alcohol to obtain *trans*18:1ω5.

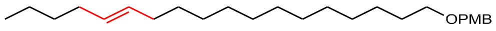


**(E)-1-methoxy-4-((octadec-13-en-1-yloxy)methyl)benzene;** Molecular Formula: $C_{26}H_{44}O_2$; Rf (30% ethyl acetate/hexanes): 0.2; **$^1$H NMR (600 MHz, $CDCl_3$)** δ 7.30 – 7.27 (m, 2H), 6.90 (d, *J* = 8.6 Hz, 2H), 5.44–5.37 (m, 2H), 4.45 (s, 2H), 3.83 (s, 3H), 3.45 (t, *J* = 6.7 Hz, 2H), 2.02–1.96

(m, 3H), 1.66 – 1.56 (m, 3H), 1.38 – 1.32 (m, 7H), 1.31–1.25 (m, 15H), 0.91 (t, $J$ = 6.9 Hz, 3H); **$^{13}$C NMR (151 MHz, CDCl$_3$)** δ 159.09, 130.86, 130.37, 130.30, 129.19, 113.75, 72.50, 70.26, 55.27, 32.61, 32.27, 31.85, 29.79, 29.67, 29.65, 29.63, 29.61, 29.53, 29.50, 29.18, 26.22, 22.19, 13.95.

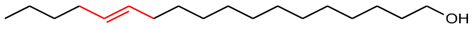


**(E)-octadec-13-en-1-ol;** Molecular Formula: $C_{18}H_{36}O$; Rf (25% ethyl acetate/hexanes): 0.2; **$^{1}$H NMR (600 MHz, CDCl$_3$)** δ 5.42 – 5.40 (m, 2H), 3.66 (t, $J$ = 6.6 Hz, 2H), 2.01 – 1.97 (m, 4H), 1.61– 1.57 (m, 3H), 1.38 – 1.27 (m, 22H), 0.91 (t, $J$ = 7.2 Hz, 3H); **$^{13}$C NMR (151 MHz, CDCl$_3$)** δ 130.36, 130.30, 63.11, 32.83, 32.60, 32.27, 29.67, 29.63, 29.62, 29.61, 29.59, 29.51, 29.43, 29.16, 25.74, 22.18, 13.94.

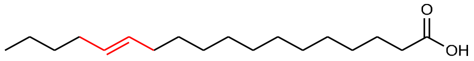


**(E)-octadec-13-enoic acid;** Molecular Formula: $C_{18}H_{34}O_2$; Rf (50% ethyl acetate/hexanes): 0.2; **$^{1}$H NMR (800 MHz, CDCl$_3$)** δ 5.39-5.38 (m, Hz, 1H), 2.35 (t, $J$ = 7.5 Hz, 1H), 2.01 – 1.93 (m, 2H), 1.65-1.61 (m, 1H), 1.37 – 1.24 (m, 17H), 0.92 – 0.86 (m, 2H); **$^{13}$C NMR (201 MHz, CDCl$_3$)** δ 178.82, 130.36, 130.31, 33.87, 32.61, 32.28, 31.84, 29.71, 29.66, 29.60, 29.58, 29.51, 29.44, 29.25, 29.16, 29.07, 24.71, 22.20, 13.97.

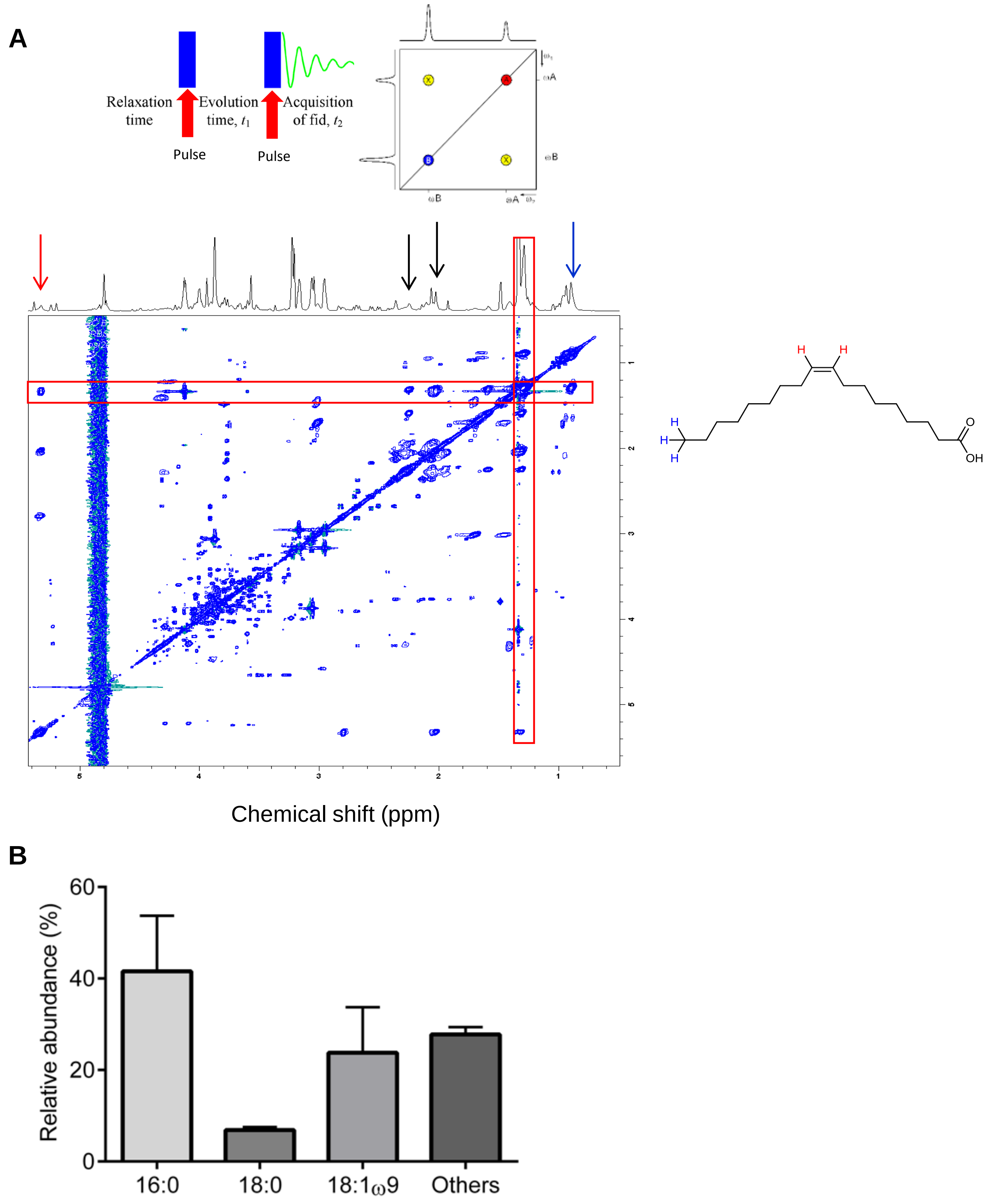


**Fig. S1** (A) Two-dimensional Correlation Spectroscopy of human neural stem and progenitor cells identifies *cis*-mono-unsaturated fatty acids, resonating at 0.88 ($CH_3$ tail, 'blue' hydrogens), 1.28 (methylene carbon backbone), 1.58, 2.02, 2.26 and 5.32ppm (*cis* formation, 'red' hydrogens). Spectra were acquired using 800MHz NMR (Bruker, Inc.; see Methods). (B) Gass Chromatography-Mass identified 18:1ω9 as the most abundant mono-unsaturated fatty acid in human neural stem and progenitor cells. Relative abundance is reported as the percent of total fatty acid methyl-esters present (N=2).

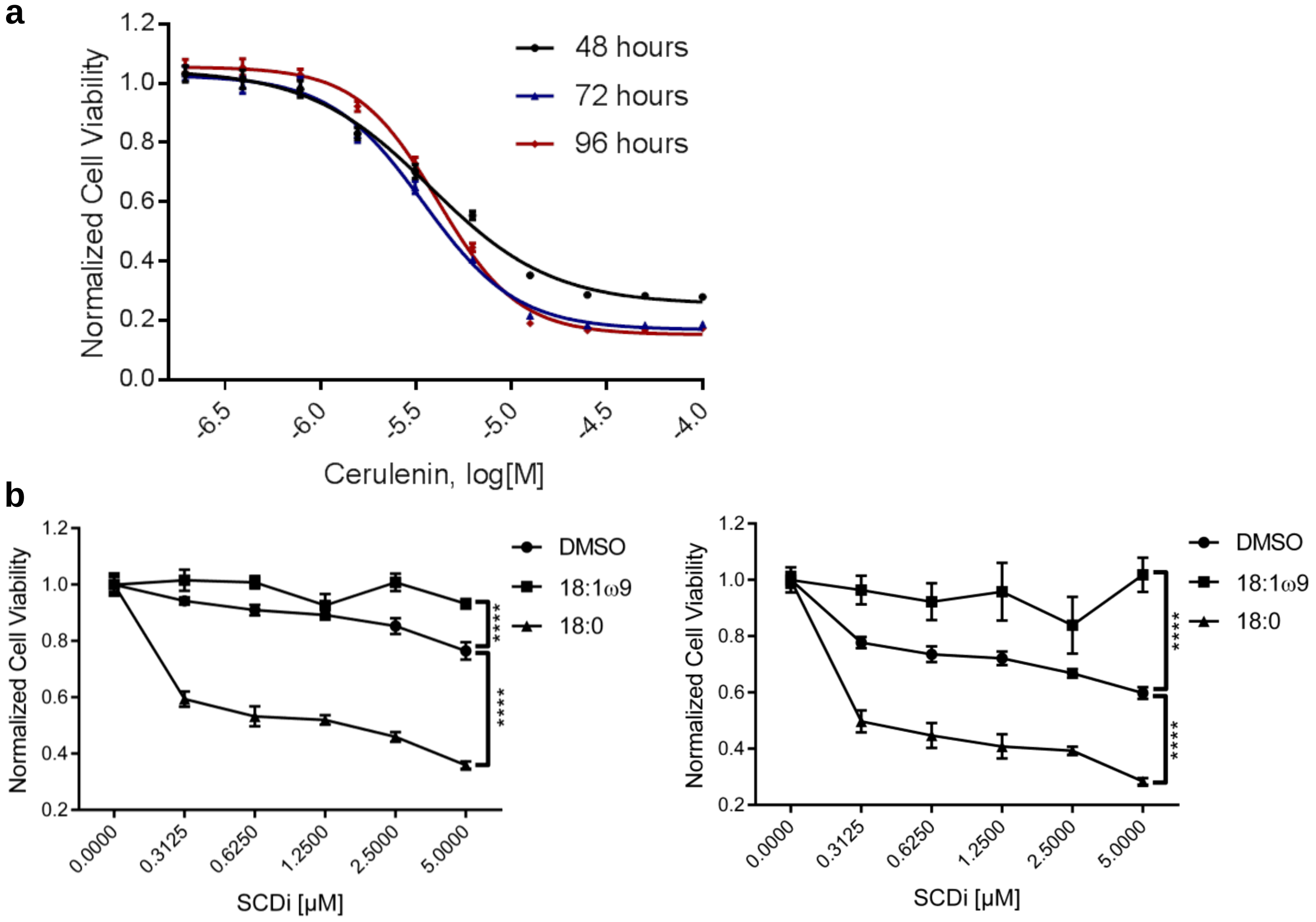


**Fig. S2 Cell growth of human neural stem and progenitor cells depends on fatty acid synthesis and production of mono-unsaturated fatty acids such as oleic acid (18:1ω9).** (A) Dose-dependent inhibition of fatty acid synthase with cerulenin reduces human neural stem and progenitor cell viability assessed by MTT assay. $EC_{50}$ was 37.5μM (95% CI= 34.9 to 40.3μM), 34.4μM (95% CI= 32.8 to 36.1μM), 41.0μM (95% CI= 39.5 to 42.6μM) at 48, 72 and 96hr, respectively. MTT absorbance was normalized to vehicle (DMSO)-treated wells (N≥3 per group). (B) 18:1ω9, but not its precursor 18:0 SFA, rescues cell viability following 72hr (*left graph*) and 96hr (*right graph*) of stearoyl-CoA desaturase inhibitor (SCDi) treatment. MTT absorbance for each timepoint was normalized to vehicle (DMSO)-treated wells (N≥3 per group); bar graphs represent mean ± SD. Two-way ANOVA test, ****p≤0.0001.

**A**

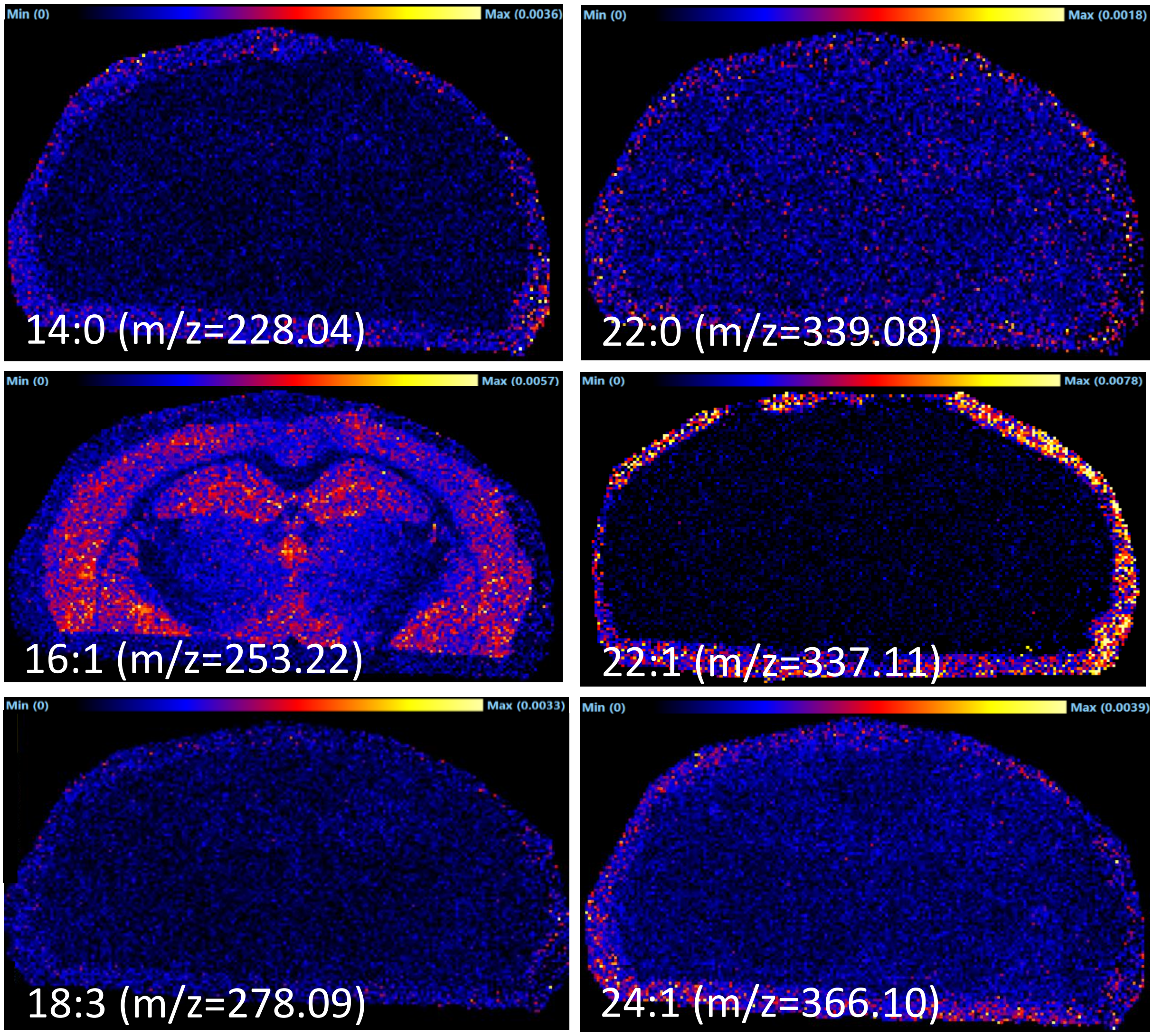


**B**

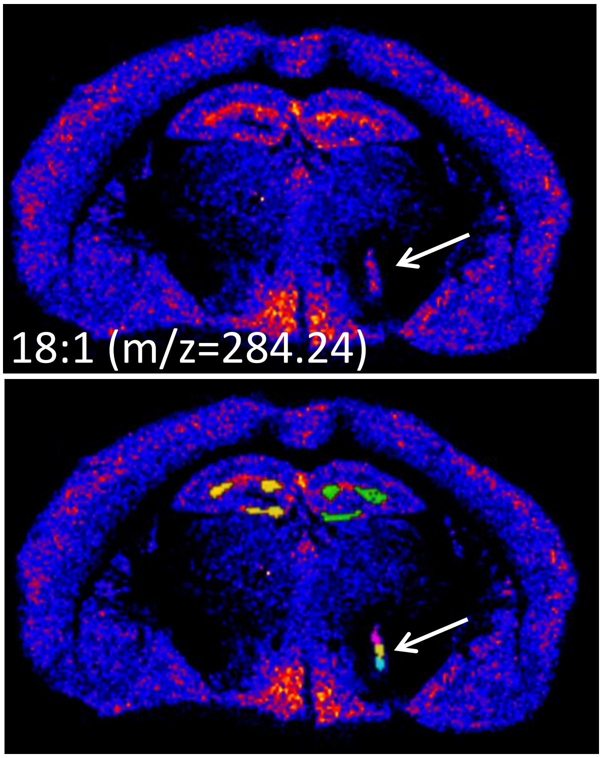


| SAMPLE | DGLEFT | DGRIGHT | SPOT |
|---|---|---|---|
| 1 | 5.163 | 5.308 | 5.416 |
| 2 | 5.096 | 5.11 | 5.215 |
| 3 | 5.253 | 5.052 | 4.939 |

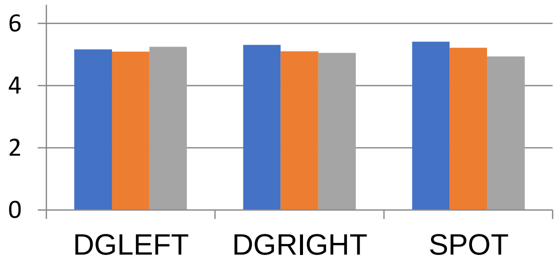


**Fig. S3** Imaging mass spectrometry of the 3-month-old wild-type C57BL/6J mouse brain using a MALDI TOF mass spectrometer (Waters, Inc; see Methods for details) shows distribution of the selected fatty acids. (A) Most fatty acids could not be detected in the mouse brain while some, such as 16:1 MUFA ion (m/z 253.22), were distributed in different brain regions, including the hippocampus. 18:3 MUFA (m/z 278.09) ion was not detected. (B) 0.5mM 18:1 MUFA was spiked-in (arrow). Three regions of the dentate gyrus (DG) as well as the spiked area (arrow) were sampled. The table and the bar graph show normalized signal intensities (Progenesis, Inc.).

**A**

| Gene ID | NCBI Accession Number | % amino acid match compared to human TLX |
|---|---|---|
| Nr2e1 | NP_689415.1 | 99% |

**B**

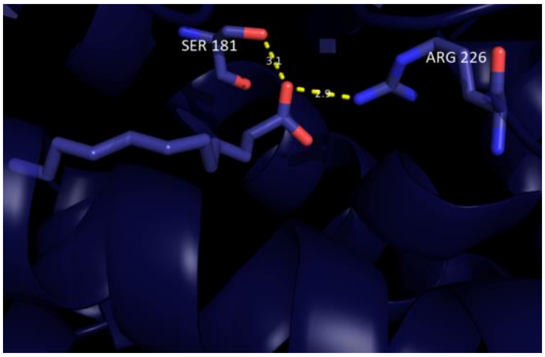


**C**

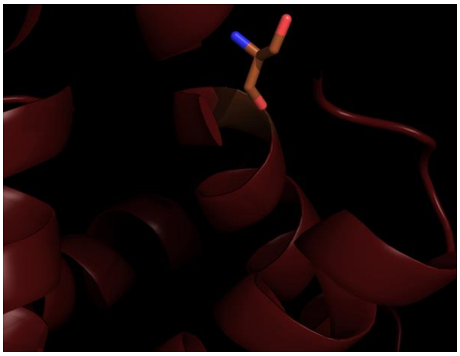

**D**

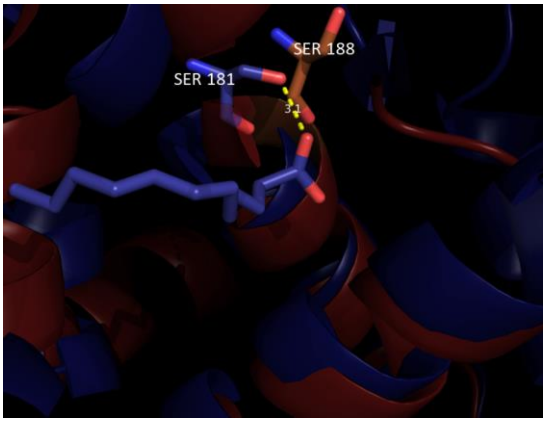


**E**

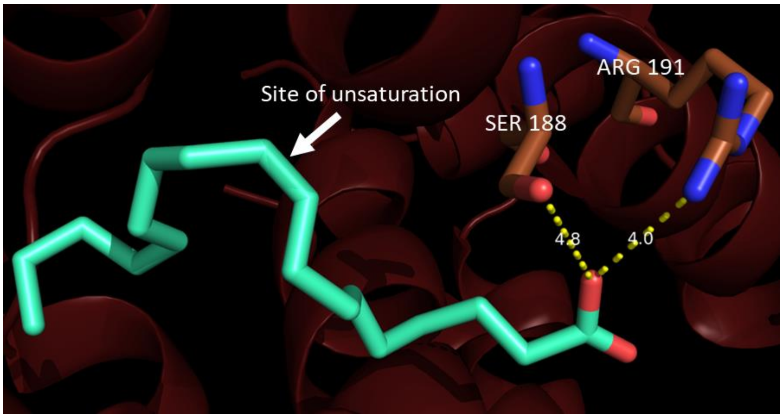


**Fig. S4 TLX ligand-binding domain homology model indicates potential fatty acid binding pocket.** (A) The full-length human TLX amino acid sequence is 99% identical to mouse Tlx, with a single residue difference in the ligand-binding domain (data not shown). Comparison was done using NCBI blastp tool. (B-E) TLX ligand-binding domain homology model based on the fatty acid-bound HNF4α and fatty acid docking experiments to TLX ligand-binding domain. (B) Protein-ligand interactions in the crystal structure of lauric acid-bound HNF4α ligand-binding domain (PDB 1M7W). (C) Conformation of TLX ligand-binding domain putative ligand-binding pocket based on the homology model generated using ligand-bound HNF4α. (D) Overlay of fatty acid-bound ligand-binding domains of HNF4α and TLX reveal that TLX may bind hydrophobic ligands such as fatty acids. (E) Molecular docking of 18:1ω9 to the putative ligand binding pocket of TLX ligand-binding domain indicates potential ligand-binding interaction.

**A**

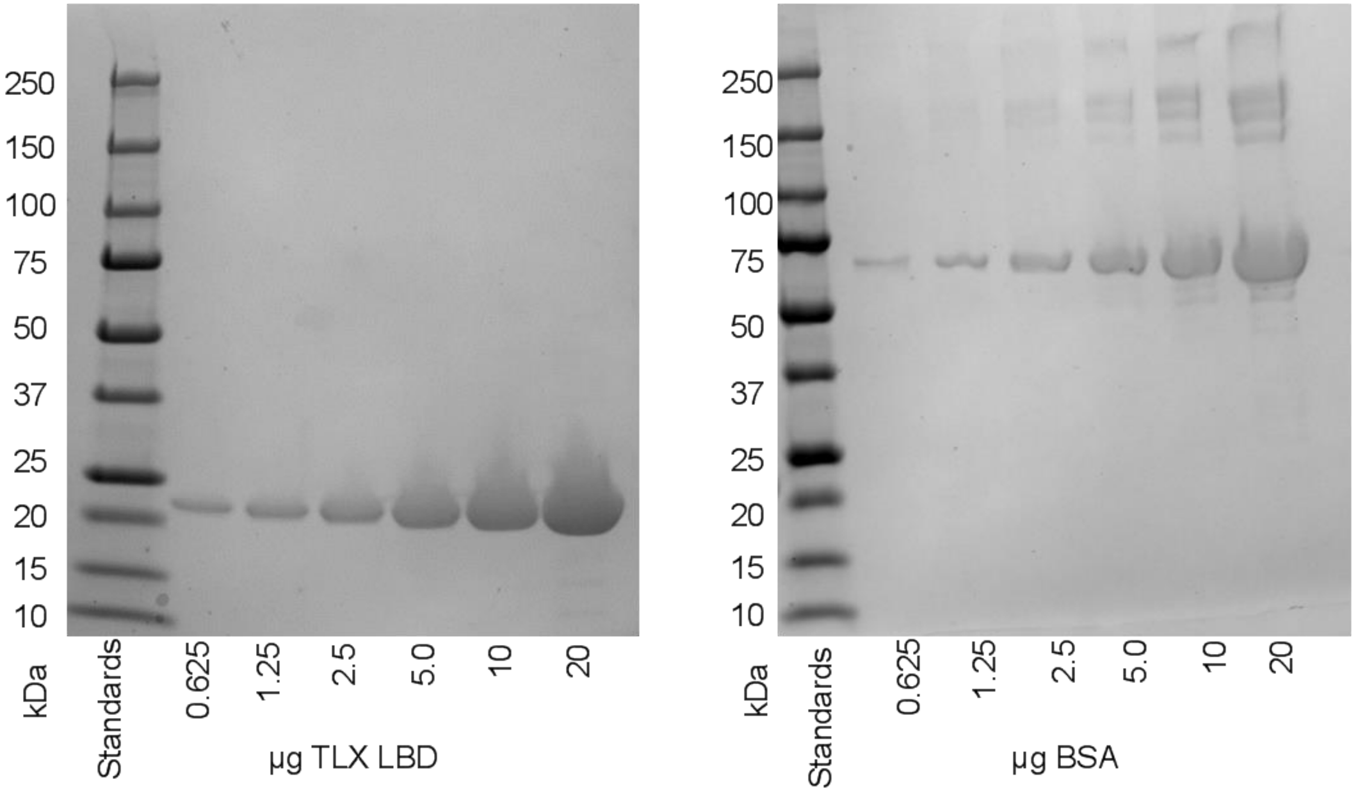


**B**

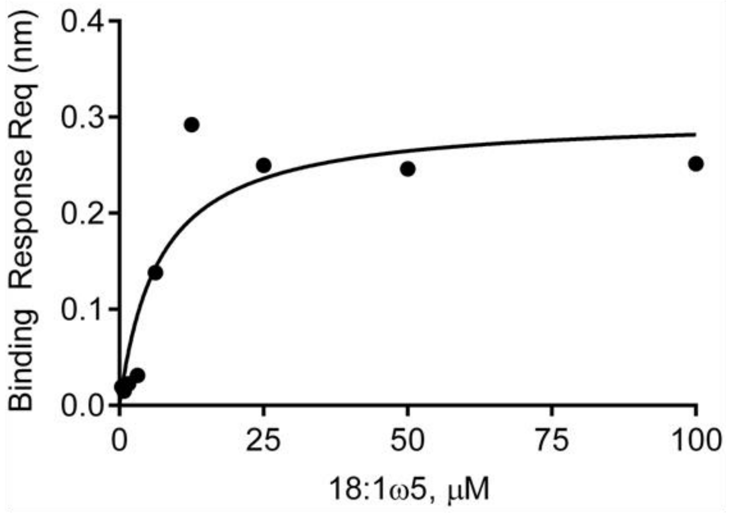


**Fig. S5 Expression and purification of TLX ligand-binding domain.** (A) Analysis of purified recombinant TLX ligand-binding domain by SDS-PAGE and Coomassie staining (*left panel*); the same amounts of bovine serum albumin (BSA) were analyzed in parallel (*right panel*). (B) Direct binding response of a synthetic MUFA, 18:1ω5, shows saturable binding response to TLX ligand-binding domain by biolayer interferometry. 18:1ω5 binds with a dissociation constant ($K_d$) of 6.5µM (95%CI=-0.2 to 13µM), calculated based on the one-site binding model.

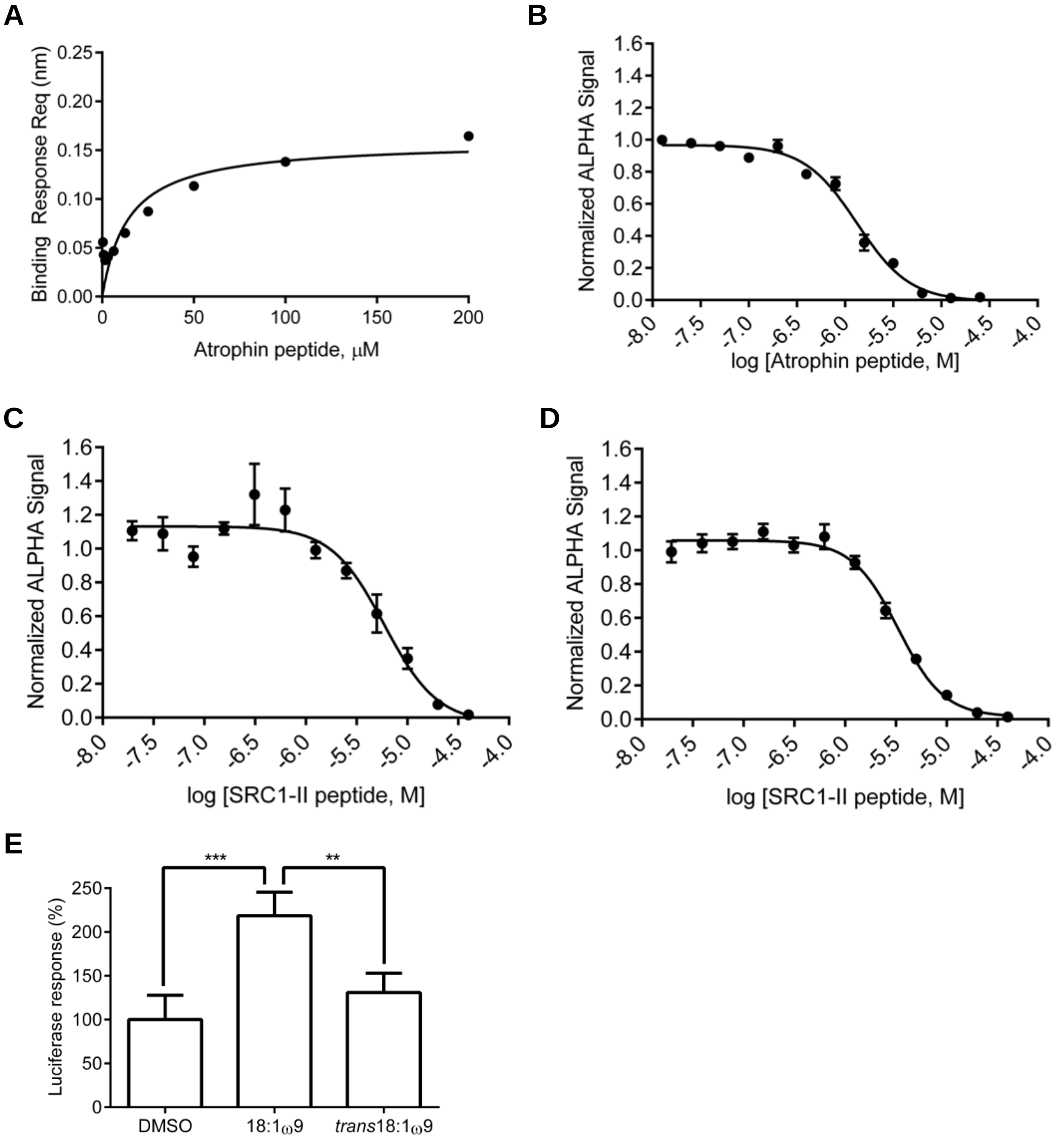


**Fig. S6 ALPHA screen-based competition assays demonstrate TLX ligand-binding domain binding to the corepressor atrophin in the absence of any ligands, and to the coactivators, nuclear receptor coactivators (NCOAs), in the presence of 18:1ω9.** (A) Atrophin peptide binds to TLX ligand-binding domain with the $K_d$ of 14.2μM (95%CI=-2.6 to 31.1μM) using biolayer interferometry assay. (B) Competition experiments with non-biotinylated atrophin peptide and TLX ligand-binding domain bound to biotinylated atrophin peptide, in the absence of fatty acid ligands. The half-maximum effective concentration ($EC_{50}$) for non-biotinylated atrophin peptide binding to TLX ligand-binding domain was 1.3μM (95%CI=1.2 to 1.5μM). (C) Competition experiments with non-biotinylated NCOA1-II peptide and TLX ligand-binding domain-bound biotinylated NCOA1-II peptide in the presence of 200μM 18:1ω9 revealed an $EC_{50}$ of 5.6μM (95% CI =4.2 to 7.7μM). (D) Competition experiments with non-biotinylated NCOA1-II peptide and biotinylated TLX ligand-binding domain-bound to GST-NCOA3-RID in the presence of 200μM 18:1ω9 revealed an $EC_{50}$ of 3.3μM (95% CI = 3.0 to 3.7μM). N≥3 for all experiments. (E) Nucleofection of TLX-based dual luciferase reporter with TLX-expressing plasmid in HeLa cells. Comparisons show increased luciferase response with 18:1ω9 but not *trans*18:1ω9 (200μM each), highlighting the conformational specificity of TLX in cells. Bar graphs represent mean ± SD. DMSO vs. *trans*18:1ω9 did not significantly differ based on Tukey's multiple comparisons test, **p≤0.01, ***p≤0.001, N≥3 for all data points.

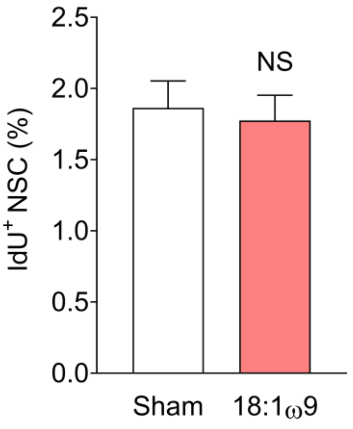


**Fig. S7. Mitotic effect of 18:1ω9 on radial neural stem cells is TLX-dependent.** To determine long-term effects of 18:1ω9 on radial neural stem cells, mice were injected with IdU (115mg/kg, 4 intraperitoneal injections 2hr apart) 30 days post-sham or fatty acid delivery (N=3-4 per group). For statistical analysis, quantitation was done using data from both dentate gyri. The percentage of proliferating radial neural stem cells did not differ from sham-injected mice 30 days post-injection of the 18:1ω9.

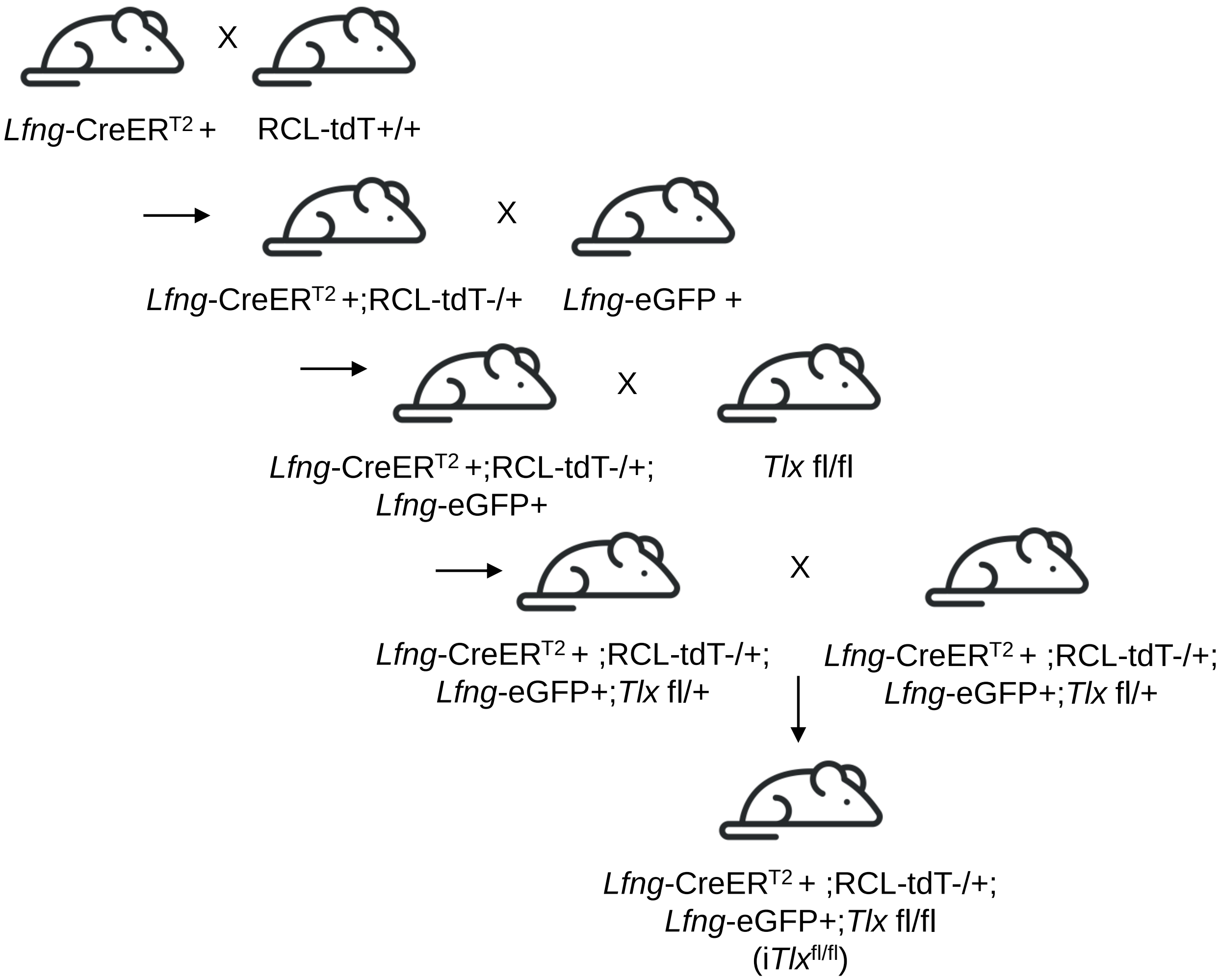


**Fig. S8 Generation of mice with selective inducible Tlx knockout in neural stem cell reporter mice**. Transgenic mouse crossings performed to generate conditional homozygous deletion of *Tlx* specifically in radial neural stem cells. Crossing with the *Lfng*-eGFP transgenic mouse ensures that all radial neural stem cells express eGFP. Mutant clones express tdTomato in addition to eGFP.

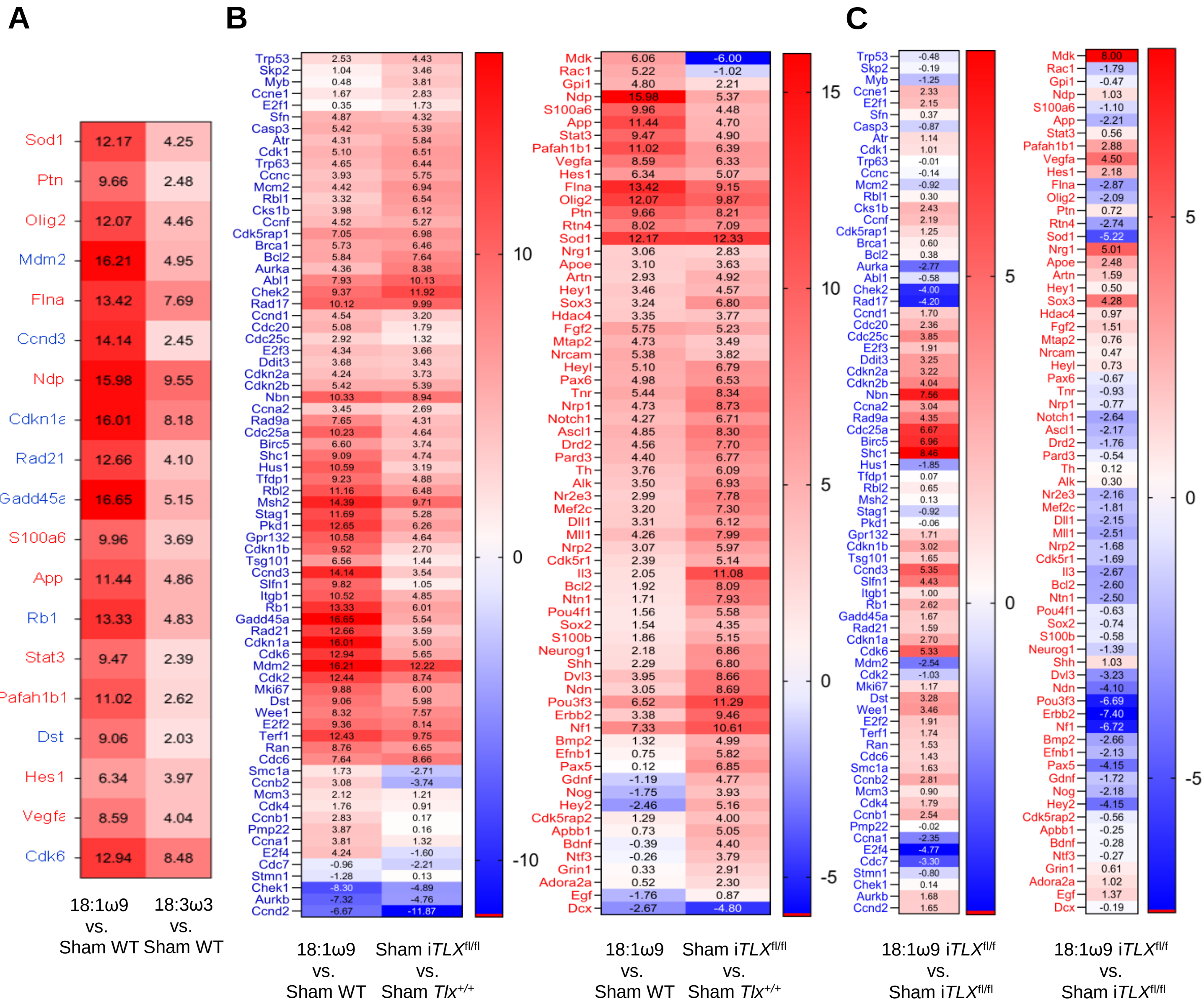


**Fig. S9** (A) Heatmap of ΔΔCt values (cited in boxes) indicates 8 cell cycle (blue letters) and 11 neurogenesis (red letters) genes upregulated by 18:3ω3 in *Lfng*-eGFP$^+$ neural stem cells (*Lfng*-eGFP mice) and thus excluded from the list of 18:1ω9-dependent TLX-regulated genes. Pairwise comparisons of 18:1ω9 vs. sham wild-type (WT) neural stem cells indicates genes upregulated by 18:1ω9. Pairwise comparisons of 18:3ω3 vs sham WT neural stem cells indicates genes upregulated by 18:3ω3. (B) Heatmaps of ΔΔCt values (cited in boxes) indicate 74 cell cycle (blue letters) and 67 neurogenesis (red letters) genes detected in all samples. Pairwise comparisons of 18:1ω9 vs. sham WT neural stem cells (*Lfng*-eGFP$^+$ neural stem cells from *Lfng*-eGFP$^+$ mice) indicates genes affected by 18:1ω9. Pairwise comparisons of sham i*Tlx*$^{fl/fl}$ neural stem cells (i*Tlx*$^{fl/fl}$ eGFP$^+$; tdT$^+$) vs. sham *Tlx*$^{+/+}$ neural stem cells indicates genes de-repressed by TLX. (C) Heatmaps of ΔΔCt values (cited in boxes) indicate 74 cell cycle (blue letters) and 67 neurogenesis (red letters) genes indicates effects of 18:1ω9 in the absence of *Tlx*, derived by 18:1ω9 i*Tlx*$^{fl/fl}$ neural stem cells (i*Tlx*$^{fl/fl}$ eGFP$^+$; tdT$^+$) vs. sham i*Tlx*$^{fl/fl}$ neural stem cells pairwise comparisons.

**A**

| | Contralateral | | Ipsilateral | |
|---|---|---|---|---|
| **NSC** | % EdU$^+$ | SEM | % EdU$^+$ | SEM |
| Sham | 1.96 | 0.33 | 1.88 | 1.17 |
| 18:3ω3 | 1.55 | 0.22 | 1.45 | 0.57 |
| 18:1ω9 | 2.37 | 0.61 | 8.97 | 1.69 |

| | Contralateral | | Ipsilateral | |
|---|---|---|---|---|
| **ANP** | % EdU$^+$ | SEM | % EdU$^+$ | SEM |
| Sham | 8.23 | 1.12 | 7.48 | 5.19 |
| 18:3ω3 | 3.78 | 0.56 | 7.56 | 2.4 |
| 18:1ω9 | 5.98 | 0.68 | 23.99 | 5.28 |

| **Two-way ANOVA** | P value |
|---|---|
| Sham vs 18:1ω9 vs 18:3ω3 | 0.0012 |
| Contralateral vs Ipsilateral | 0.0281 |

| **Two-way ANOVA** | P value |
|---|---|
| Sham vs 18:1ω9 vs 18:3ω3 | 0.0112 |
| Contralateral vs Ipsilateral | 0.0092 |

| **Tukey's Multiple Comparison Test** | |
|---|---|
| Contralateral | Ipsilateral |
| Sham vs 18:3ω3=0.97 | Sham vs 18:3ω3=0.95 |
| Sham vs 18:1ω9=0.97 | Sham vs 18:1ω9=0.0002 |
| 18:3ω3 vs 18:1ω9=0.89 | 18:3ω3 vs 18:1ω9 <0.0001 |

| **Tukey's Multiple Comparison Test** | |
|---|---|
| Contralateral | Ipsilateral |
| Sham vs 18:3ω3=0.59 | Sham vs 18:3ω3=1.0 |
| Sham vs 18:1ω9=0.85 | Sham vs 18:1ω9=0.0028 |
| 18:3ω3 vs 18:1ω9=0.87 | 18:3ω3 vs 18:1ω9=0.0013 |

**B**

| | ***Tlx*$^{+/+}$** | | | | **i*Tlx*$^{fl/+}$** | | | | **i*Tlx*$^{fl/fl}$** | | | |
|---|---|---|---|---|---|---|---|---|---|---|---|---|
| | Contralateral | | Ipsilateral | | Contralateral | | Ipsilateral | | Contralateral | | Ipsilateral | |
| **NSC** | % EdU$^+$ | SEM | % EdU$^+$ | SEM | % EdU$^+$ | SEM | % EdU$^+$ | SEM | % EdU$^+$ | SEM | % EdU$^+$ | SEM |
| Sham | 1.58 | 0.37 | 1.82 | 0.49 | 0.93 | 0.56 | 0.82 | 0.51 | 0.62 | 0.45 | 0.92 | 0.63 |
| 18:1ω9 | 2.42 | 0.68 | 9.97 | 1.12 | 1.8 | 0.35 | 5.69 | 0.63 | 0.68 | 0.49 | 0.57 | 0.37 |

| **Two-way ANOVA** | ***Tlx*$^{+/+}$** | **i*Tlx*$^{fl/+}$** | **i*Tlx*$^{fl/fl}$** |
|---|---|---|---|
| | P value | P value | P value |
| Sham vs 18:1ω9 | <0.0001 | 0.0004 | 0.7868 |
| Contralateral vs Ipsilateral | <0.0001 | 0.0102 | 0.8562 |

| **Tukey's Multiple Comparison Test** | ***Tlx*$^{+/+}$** | **i*Tlx*$^{fl/+}$** | **i*Tlx*$^{fl/fl}$** |
|---|---|---|---|
| Sham:Contralateral vs. Sham:Ipsilateral | 0.9966 | 0.9997 | 0.9803 |
| Sham:Contralateral vs. 18:1ω9:Contralateral | 0.87 | 0.7873 | 0.9998 |
| Sham:Contralateral vs. 18:1ω9:Ipsilateral | <0.0001 | 0.0004 | >0.9999 |
| Sham:Ipsilateral vs. 18:1ω9:Contralateral | 0.9474 | 0.7256 | 0.9868 |
| Sham:Ipsilateral vs. 18:1ω9:Ipsilateral | <0.0001 | 0.0003 | 0.9511 |
| 18:1ω9:Contralateral vs. 18:1ω9:Ipsilateral | <0.0001 | 0.0002 | 0.9981 |

**Table 1. Mitotic effect of 18:1ω9 on radial neural stem cells (NSCs) is TLX-dependent**. (A) The effects of either sham, 18:1ω9, or 18:3ω3 on radial NSC and amplifying neuroprogenitor (ANP) proliferation (EdU+ cells). Statistical analyses related to **Fig. 3C.** (B) The effects of either sham or 18:1ω9 on radial NSC proliferation (EdU+ cells) in mice lacking *Tlx*. Statistical analyses related to **Fig. 3E.**

| | Sham | | 18:3ω3 | | *cis*18:1ω9 | |
|---|---|---|---|---|---|---|
| | # of $CldU^+$ cells | SEM | # of $CldU^+$ cells | SEM | # of $CldU^+$ cells | SEM |
| NeuN | 600.25 | 42.64 | 582.24 | 59.54 | 1065 | 52.84 |
| Dcx | 73.63 | 25.01 | 106.6 | 1.13 | 449.24 | 37.4 |

| $NeuN^+$ $CldU^+$ | |
|---|---|
| Two-way ANOVA | P value |
| Sham vs. 18:3ω3 | 0.9622 |
| Sham vs. *cis*18:1ω9 | <0.0001 |
| 18:3ω3 vs. *cis*18:1ω9 | <0.0001 |

| $Dcx^+$ $CldU^+$ | |
|---|---|
| Two-way ANOVA | P value |
| Sham vs. 18:3ω3 | 0.8794 |
| Sham vs. *cis*18:1ω9 | <0.0001 |
| 18:3ω3 vs. *cis*18:1ω9 | <0.0001 |

**Table 2. 18:1ω9 increases neurogenesis *in vivo.*** The effects of either sham, 18:1ω9, or 18:3ω3 on the number of newborn immature neuron ($CldU^+$ $Dcx^+$) and granule cells ($CldU^+$ $NeuN^+$).
